# Extracellular vesicle-mediated trafficking of developmental cues is altered during human brain disease

**DOI:** 10.1101/2023.06.27.546646

**Authors:** Fabrizia Pipicelli, Andrea Forero, Sylvain Moser, Francesco Di Matteo, Natalia Baumann, Christian Grätz, Mariano Gonzalez Pisfil, Zagorka Bekjarova, Michael W. Pfaffl, Laura Canafoglia, Benno Pütz, Pavel Kielkowski, Filippo M. Cernilogar, Giuseppina Maccarrone, Denis Jabaudon, Rossella Di Giaimo, Silvia Cappello

**Affiliations:** Max Planck Institute of Psychiatry, Munich, Germany; International Max Planck Research School for Translational Psychiatry; Department of Basic Neurosciences, University of Geneva, Geneva, Switzerland; Division of Animal Physiology and Immunology, Technical University of Munich, Freising, Germany; Core Facility Bioimaging and Walter-Brendel-Centre of Experimental Medicine, Biomedical Center, Ludwig-Maximilians-University, Munich, Germany; Fondazione IRCCS Istituto Neurologico Carlo Besta, Milan, Italy; Department of Chemistry, Ludwig-Maximilians-University, Munich, Germany; Biomedical Center, Faculty of Medicine, Ludwig-Maximilians-University, Munich, Germany; Department of Biology, University of Naples Federico II, Naples, Italy

**Keywords:** extracellular vesicle, epilepsy, neurodevelopment, cerebral organoids

## Abstract

Cellular crosstalk is an essential process influenced by numerous factors including secreted vesicles that transfer nucleic acids, lipids, and proteins between cells. Extracellular vesicles (EVs) have been the center of many studies focusing on neuron-to-neuron communication, but the role of EVs in progenitor-to-neuron and -astrocyte communication and whether EVs display cell-type-specific features for cellular crosstalk during neurogenesis is unknown. Here, using human-derived cerebral organoids, neural progenitors, neurons, and astrocytes, we found that many proteins coded by genes associated with neurodevelopmental disorders are transported via EVs. Thus, we characterized the protein content of EVs and showed their cell type-specific dynamics and function during brain development. Changes in the physiological crosstalk between cells can lead to neurodevelopmental disorders. EVs from patients with epilepsy were found altered in composition and function. Alterations in the intracellular and extracellular compartments highlighted a clear dysregulation of protein trafficking. This study sheds new light on the biology of EVs during brain development and neurodevelopmental disorders.

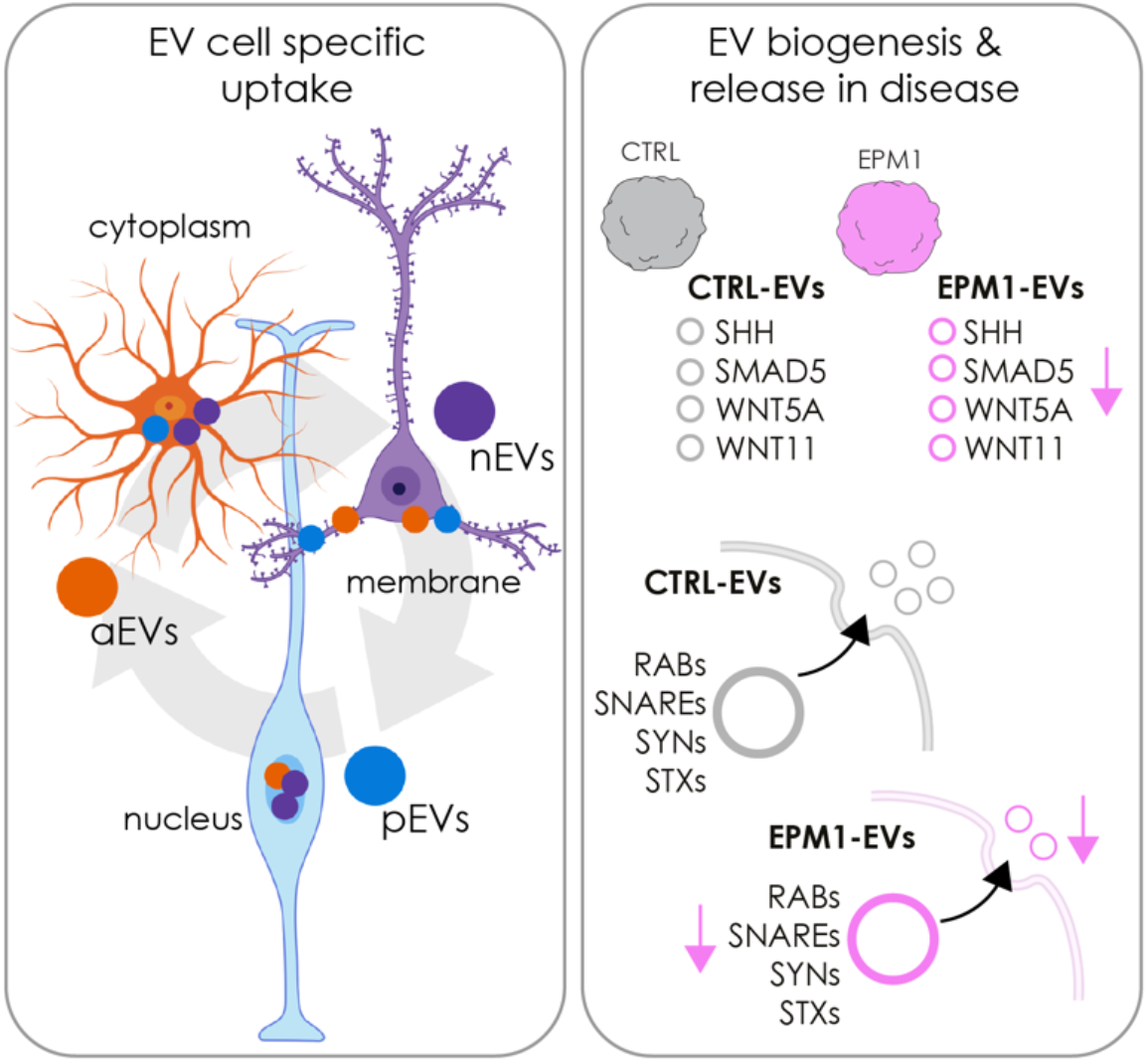

**Graphical abstract:** (left) EV uptake mechanism varies depending on the receiving cell type; NPCs transport neuron EVs (nEVs) and astrocyte EVs (aEVs) to the nucleus, astrocytes localize progenitor EVs (pEVs) to the cytoplasm, and neurons retain pEVs and aEVs along the plasma membrane. (right) Cerebral organoids (COs) from progressive Myoclonus Epilepsy Type I (EPM1) patients release EVs lacking key proteins in neurodevelopment and proteins necessary for EV biogenesis and release. Illustration created using BioRender.

## Introduction

The events that organize brain structure during development include neurogenesis, cell migration as well as axon projection and guidance (Heng et al., 2010; Rakic, 2009; Taverna et al., 2007). In this dynamic context, cell-to-cell communication is an essential process influenced by factors, including cell morphology, adhesion molecules, the local extracellular matrix (ECM) and secreted vesicles (Long et al., 2018; Pellegrini et al., 2020; Peruzzotti-Jametti et al., 2021). Extracellular signals are required during development to establish precise cell numbers, unique cell types, and specific cell migration patterns, positions, and functions (Long and Huttner, 2019; Sharma et al., 2019; Silva et al., 2019).

Extracellular vesicles (EVs) are small particles in the brain extracellular environment which have gained interest due to their potential in diagnostics and therapeutics in neurodevelopmental disorders (Gomes et al., 2020). EVs are secreted by all cells and are classified mostly as exosomes and microvesicles based on their size, composition, and origin. While exosomes are generally smaller (30-200 nm) and released by fusion of multivesicular bodies with the plasma membrane, microvesicles (of much variable size, 100-1000 nm) are directly shed by the outward blebbing of the plasma membrane (Van Niel et al., 2018). Vesicles travel long distances within and outside of cells, thus impacting crosstalk at several levels and sites. Although EVs are key players in extracellular environment composition, few studies have focused on their functional role during brain development and disease. Neural stem cells secrete EVs containing different cargoes including Prominin-1 (Marzesco et al., 2005). Similarly, retinal stem cells release EVs containing developmental transcription factors, microRNA and membrane proteins that regulate gene expression in developing mouse and human retinal organoids (Zhou et al., 2018, 2021). Neurons and astrocytes in vitro have also been shown to secrete EVs containing varied cargoes, such as L1 adhesion molecule and specific subunits of the glutamate receptor (Fauré et al., 2006). EVs also have physiological and pathological functions, for instance directing neuronal differentiation and regulating synapse formation in healthy neurons (Fauré et al., 2006; Schiera et al., 2015; Takeda and Xu, 2015) as well as in neurodevelopmental disorders such as cortical malformations and autism spectrum disorder (Kyrousi et al., 2021; Pipicelli et al., 2023; Sharma et al., 2019).

Recent studies support that EVs, comprising both exosomes and microvesicles, can act as carriers of ECM components such as Tenascin C (Albacete-Albacete, 2021; Rilla et al., 2019), and regulate cell growth, differentiation and cell migration by transporting ECM remodeling cargoes like matrix metalloproteinases that can alter depositions of different collagen types (Nawaz et al., 2018). Moreover, annexin-enriched osteoblast-derived EVs act as an extracellular site of mineral nucleation in developing stem cell cultures (Davies et al., 2017).

During brain development, ECM alterations and secreted factors can lead to neuronal disfunction (Long and Huttner, 2019) and therefore contribute to the occurrence of neurodevelopmental disorders. In previous work, we present how Cystatin B (CSTB), a ubiquitous protease inhibitor whose mutation is responsible for progressive Myoclonus Epilepsy Type I (EPM1), is secreted and has an essential role in instructing neighboring cells during neurogenesis (Di Matteo et al., 2020). We hypothesize the secretion of CSTB via EVs as a new mechanism of extracellular environment modification. It is therefore challenging but crucial to decode how EV signals are coordinated during human brain development and if altered EV crosstalk can impact the correct establishment of the brain architecture and lead to neurodevelopmental disorders.

Here we present insight into EV release, composition, uptake, and function using human brain models: cells in 2D monolayer and a 3D model of human brain development, namely cerebral organoids (COs). Additionally, we present a systematic analysis of EV content in association with some neurodevelopmental disorders, and more specifically, EPM1.

## Results & Discussion

### EV Isolation and cell type-specific characterization

We isolated a mixed population of EVs containing both exosomes and small microvesicles (100-300 nm) from the secreted fraction (culture medium) of COs and 2D cell cultures due to the challenges of isolating specific subtypes of EVs, since exosomes and microvesicles share common markers and sizes. EV isolation was done by differential ultracentrifugation (Mathieu et al., 2019; Théry et al., 2006) and we further characterized EVs by nanoparticle tracking analysis (NTA) and immuno-electron microscopy with transmembrane (CD81) and intraluminal (LGALS3) markers (Fig. 1A-B and Fig. S1A).

**Fig. 1.**
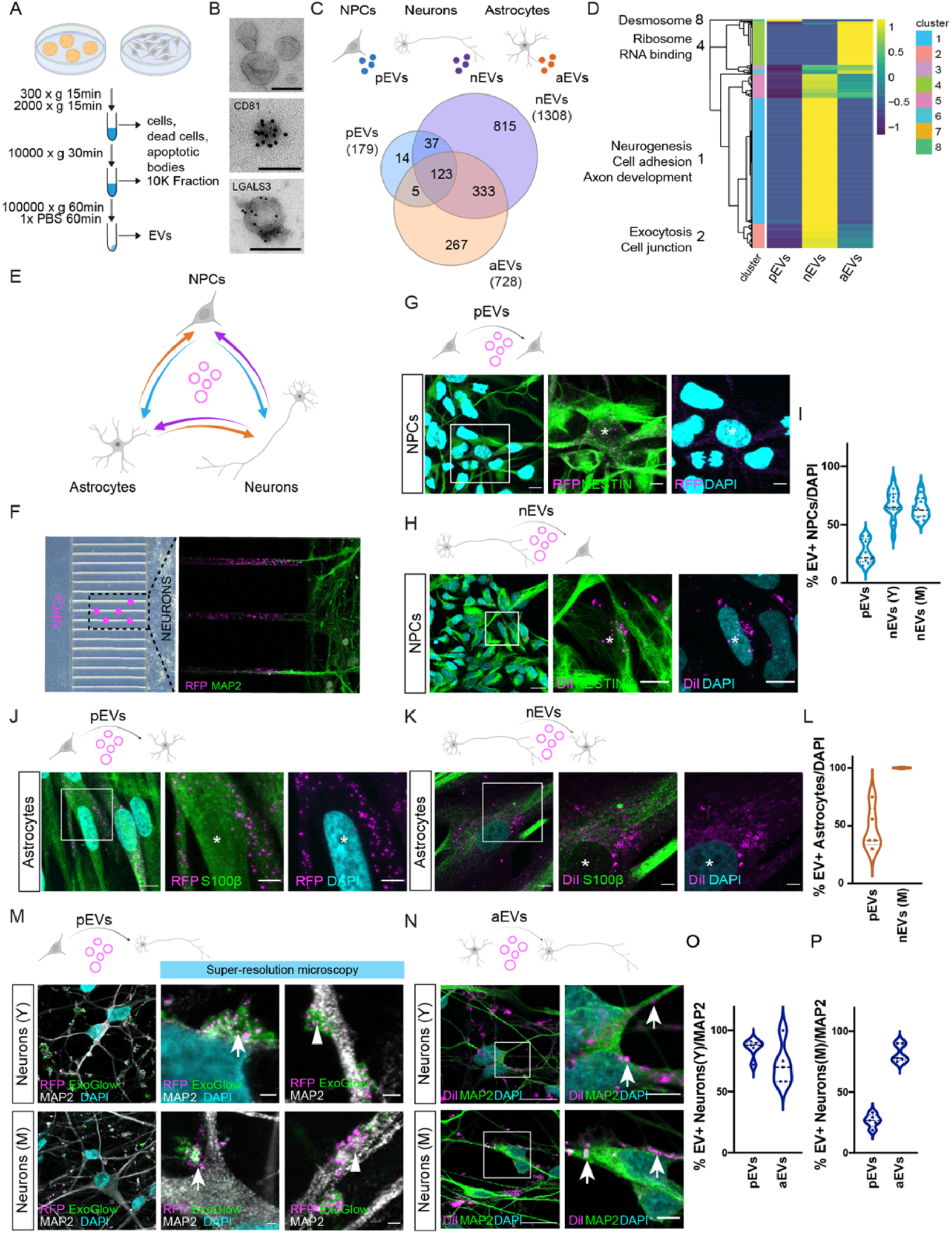
Isolation and cell-type-specific characterization of EVs. (A) Schematic of EV isolation protocol by differential ultracentrifugation. (B) Immuno-electron micrographs of CD81 and LGALS3 in EVs collected from COs conditioned media. Scale bar:100nm. (C) Schematic of EVs secreted by 2D cell populations: NPCs (neural progenitor cells, pEVs, blue), neurons (nEVs, purple) and astrocytes (aEVs, orange) (top), and Venn diagram of proteins in pEVs, nEVs, and aEVs (bottom). (D) Heatmap showing hierarchical clusters of proteins in pEVs, nEVs, and aEVs. GO enrichments of clusters are displayed. EVs were collected from the conditioned media of 3-5 different wells of cells in culture. (E) Schematic of the experimental setup of EV exchange between different cell types. (F) Microfluidic chamber showing the stream of EVs (RFP+) secreted by NPCs traveling to neurons (MAP2+). (G) Immunostaining indicating uptake of pEVs (magenta) by NPCs (Nestin+). Asterisks indicate EV receiving cells. DAPI, cyan. Scale bars: full image, 10 μm; close-up 5 μm. (H) Immunostaining indicating uptake of nEVs (magenta) by NPCs (Nestin+). Asterisks indicate EV receiving cells. DAPI, cyan. Scale bars: full image, 10 μm; close-up 5 μm. (I) Quantification of EV+ NPCs following treatment with pEVs, nEVs (Y, young), nEVs (M, mature). Data are represented as mean and ± SEM. Every dot refers to a field of view, n= 9 (pEVs), n=15 (nEVs, M), n=15 (nEVs, Y). (J) Immunostaining indicating uptake of pEVs (magenta) by astrocytes (S100b+). Asterisks indicate EV receiving cells. DAPI, cyan. Scale bars: full image, 10 μm; close-up 5 μm. (K) Immunostaining indicating uptake of nEVs (magenta) by astrocytes (S100b+). Asterisks indicate EV receiving cells. DAPI, cyan. Scale bars: full image, 10 μm; close-up 5 μm. (L) Quantification of EV+ astrocytes following treatment with pEVs, nEVs (Y). Data are represented as mean and ± SEM. Every dot refers to a field of view, n= 5 (pEVs), n= 5 (nEVs). (M) Immunostaining indicating uptake of pEVs (EVs, magenta; ExoGlow, green) by young (Y, top) and mature (M, bottom) neurons (MAP2+). Super-resolution microscopy images (right) show EV localization on cell soma (arrows) and dendrites (arrow heads). DAPI, cyan. Scale bars: full image, 10 μm; close-up 1 μm. (N) Immunostaining indicating uptake of aEVs (magenta) by young (Y, top) and mature (M, bottom) neurons (MAP2+). Arrows indicate EV localization on cell soma. DAPI, cyan. Scale bars: full image, 10 μm; close-up 1 μm. (O) Quantification of EV+ Neurons(Y) following treatment with pEVs and aEVs. Data are represented as mean and ± SEM. Every dot refers to a field of view, n= 5 (pEVs), n=5 (aEVs). (P) Quantification of EV+ Neurons(M) following treatment with pEVs and aEVs. Data are represented as mean and ± SEM. Every dot refers to a field of view, n= 5 (pEVs), n=5 (aEVs). Illustrations created using BioRender.

We hypothesized that different cell types secrete vesicles enriched with cell-type-specific proteins. For this purpose, we isolated cell-type-specific EVs and profiled their protein content by Mass Spectrometry by culturing neural progenitor cells (NPCs), neurons and astrocytes in 2D monolayer cultures (Fig. 1C-D, S1B). The results of the protein profiling validated our EV purification protocol, in accordance with previous studies (Théry et al., 2018). Standard positive (CD63, TSG101 and PDCD6IP) EV markers are present in all our samples and negative (CYC1 and GOLGA2) markers are undetectable (Fig. S1C). The relative abundance of EV proteins was confirmed by WB on independent samples (Fig. S1D).

Our results showed that the three populations of cells grown in 2D share less than 8% of the total EV proteins (Fig. 1C). EVs from NPCs (pEVs) were the least diverse (in total protein number) and less than 1% of the proteins was unique for NPCs (cluster 8, Fig. 1D and S1E). On the contrary, neurons exhibited the most unique protein content in EVs (nEVs, clusters 1 and 2 in Fig. 1D and S1E) suggesting that neurons make extensive use of EVs for cellular crosstalk, as previously shown (Men et al., 2019). RNA catabolic process and ribonucleoprotein complex (cluster 4, Fig. 1D) were enriched in astrocyte EVs (aEVs). Cell-type-specific proteomic analysis showed that EVs released from different cell-types were loaded with varying amounts of EV markers (Fig. S1C), suggesting heterogeneity of EVs.

To investigate the origin of the difference in EV protein content, we performed immunohistochemical analysis of EV specific markers in different cell types in 2D. We observed some markers being ubiquitously expressed (CD81 and PDCD6IP), while others presented a more restrictive pattern of expression among different cell types.

For instance, CD82+ and CD9+ EVs were the least abundant overall, limited to a small number of NPCs and astrocytes while absent in neurons (Fig. S2A-G). In line with the hypothesis of cell-type-specific EVs, the expression of EV markers also varied among cell types as shown by single-cell RNA-Seq analysis of COs (Fig. S2H-I). Together, our data show that varying combinations and numbers of EV markers are present in different cell-types suggesting cell-specific types of EVs.

We then investigated whether EVs have a distinct function depending on receiving cells. To assess this, we collected and fluorescently labeled EVs from the three different cell populations previously described (pEVs, nEVs and aEVs, Fig. 1E). Because RFP-labeled NPCs released RFP+pEVs that travelled to MAP2+ neurons in a microfluidic chamber (Fig. 1F), we applied RFP+pEVs directly to NPCs, neurons (young and mature) and astrocytes, and observed that the uptake mechanism varied between cell-types (Fig. S2J). NPCs displayed a preference for nEV uptake, both from young (4 weeks in culture, Y) and mature (10 weeks in culture, M) neurons, compared to pEVs (Fig. 1G-I). Astrocytes appeared to also internalize both pEVs and nEVs, with a higher uptake for nEVs (Fig. 1J-L). Interestingly, while young neurons uptake both pEVs and aEVs indistinctly, mature neurons preferentially uptake EVs released by astrocytes (Fig. 1M-P). Therefore, EVs generated by the same cell type show a different uptake in recipient cell types. For example, pEVs preferentially target young neurons, their physiological partners during development (Fig. S2J). In addition to the differences in the receiving capacity, we observed differences in EV uptake localization amongst cell types. In both young and mature neurons, pEVs and aEVs (Fig. 1M-N) remained primarily docked on the cell membrane of the soma and dendrites, suggesting receptor-mediated signaling. Meanwhile, NPCs and astrocytes exhibited EV internalization, with EVs localizing also in the nuclei of NPCs (Fig. 1G-H and J-K).

Together these data indicate that certain EVs, and naturally their content, could potentially have a higher impact on a specific cell type and that the mechanism by which EVs interact with receiving cells also varies.

### Developmental characterization of EVs

Because EV crosstalk can be mediated by specific cell populations during brain development, we performed a systematic proteomic analysis of EVs at different developmental stages in COs (d15 to d360; Fig. 2A). This analysis was conducted on different batches of organoids derived from a single control cell line, since our aim was to observe changes in EV protein composition at different time points and to overcome genetic background variability. We detected a total of 3791 proteins, with substantial heterogeneity (number of detected proteins) (Fig. 2B-C and validation by WB in Fig. S3A). The total number of EV proteins from 3D COs was strongly increased compared to 2D cultures suggesting a higher variety in 3D (Fig. 2B). Accordingly, when comparing EVs from NPCs in 2D and 3D d15 COs or mature neurons/astrocytes and 3D d360 COs, the number of proteins varied greatly, suggesting that the 3D environment contributes to EV secretion as indicated from the enriched GO terms (Fig. 2D). The unique protein content for each developmental stage is associated with cell-cycle and RNA-splicing (d15), intracellular transport and mitochondrial membrane (d40), ribosome biogenesis and mitochondrion (d200) and locomotion, secretion, neuron part and cell motility (d360) (Fig. 2B-C). 6.8% of EV proteins were shared across all developmental stages, including those involved in cell junctions and secretory functions (Fig. 2B-C). EV protein difference across development suggests unique EV signatures and variety. To assess if secretion of proteins in EVs followed their physiological pattern of expression in cells, development-associated proteins that were identified in EVs were compared with their cellular gene expression (Cruceanu et al., 2022) and localization in intracellular vesicles (IVs). Surprisingly, EV proteins did not match cellular expression strictly and their trajectories did not always follow cellular compartmentalization in intracellular vesicles (IV) (Fig. 2E and S3B-C). Markers for apical radial glia, like VIM and FABP7, differed in abundance, with a peak of expression at d100 (VIM) and d200 (FABP7) (Fig. 2E). Typical markers for basal radial glia, appearing around 50d in COs, were enriched at different stages in EVs with PTPRZ1 peaking at d100 and GNG5 at d200. Early neuronal markers were detected in EVs and while DCX peaked at d15, RELN was strongly enriched at d40 (Fig. 2E). Mature neuronal markers also exhibited distinctive patterns; TUBB3 was persistently secreted in EVs while MAP2 only after 200 days. On the contrary, some of the transcription factors (TFs) typically expressed in progenitors and neurons during development were not detected in EVs (PAX6, EOMES, HOPX, Fig. 2E and S3B-C). We next examined typical EV markers (Fig. 2F) and identified unique developmental expression trajectories suggesting EV heterogeneity. Specific exosome markers (EEA1, RAB27A, RAB5B and RAB7A) or microvesicle markers (ANXA1, ANXA5, CAV1 and IMMT) (D’Acunzo et al., 2021) also displayed a different pattern of secretion during development, suggesting a time-or cell-type-regulated secretion (Fig. 2F).

**Fig. 2.**
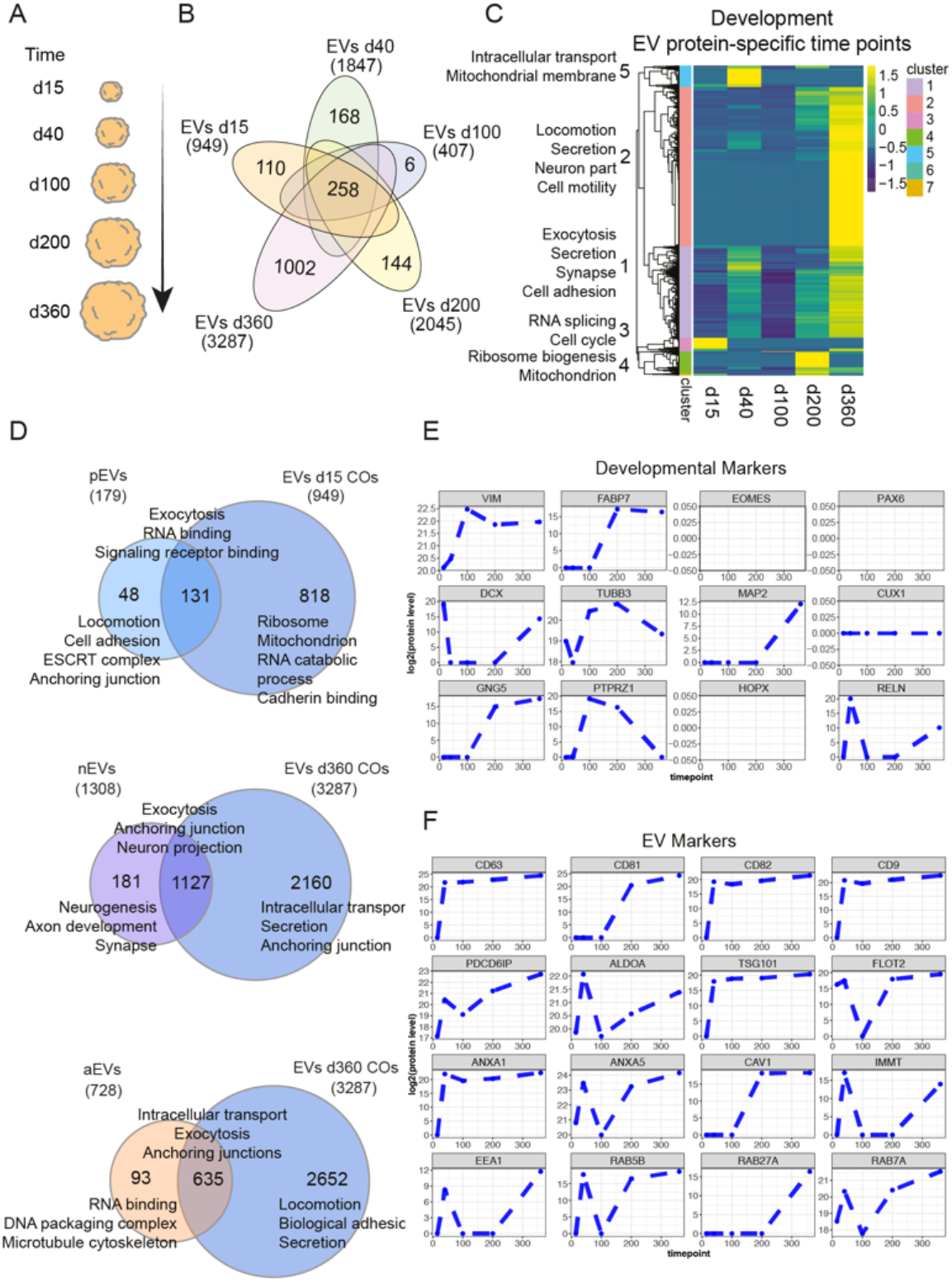
Developmental characterization of EVs. (A) Schematic of analyzed developmental stages in COs. Illustration created using BioRender. (B) Venn diagram of EV proteins at all developmental stages. (C) Heatmap showing hierarchical clusters of EV proteins at different timepoints. GO enrichments of clusters are displayed. EVs were collected from the conditioned media of 15cm petri dishes containing 20-30 COs; EVs from different timepoints were collected from at least 3 different batches of COs. (D) Venn diagram indicating the number of unique and common proteins secreted by NPCs and 15d-COs (top), neurons and 360d-COs (middle), and astrocytes and 360d-COs (bottom). Functional annotations of GO enrichments of each protein group are shown. (E) Temporal trajectories of the expression of developmental markers in EVs at different stages. (F) Temporal trajectories of EV marker expression in EVs at different stages.

Together, our results show a dynamic change in the protein content and secretion of EVs depending on the developmental stage and cell type. Moreover, a more complex environment (3D) is associated with increased EV heterogeneity.

### Brain-region-dependent signaling function of Evs

Intrigued by the idea that EV content is dependent on cell-type and context, we investigated whether EVs released from different donor cells may have a direct “signaling” function on recipient cells. Knowing that 3D organoids are a more suitable model to generate heterogeneity and diversity of EVs, we hypothesized that brain-region-specific EVs could represent different sources of EVs. Thus, we profiled EVs from dorsally- and ventrally-patterned forebrain COs (dCOs and vCOs) (Bagley et al., 2017; Birey et al., 2017)(Fig. 3A and validation by WB in Fig. S4A). EVs from dorsal (dEVs) and ventral (vEVs) COs shared 62,5% of the total proteins. While vEVs only had a small fraction of unique proteins (2.5%), dEVs contained 35% of unique proteins, showing a greater heterogeneity and suggesting that dorsal and ventral cells make distinct use of EV-mediated communication (Fig. 3A-B). Cell adhesion and motility proteins were enriched in vEVs while RNA, miRNA and chromatin binding were the main functions for dEV proteins (Fig. 3B).

**Fig. 3.**
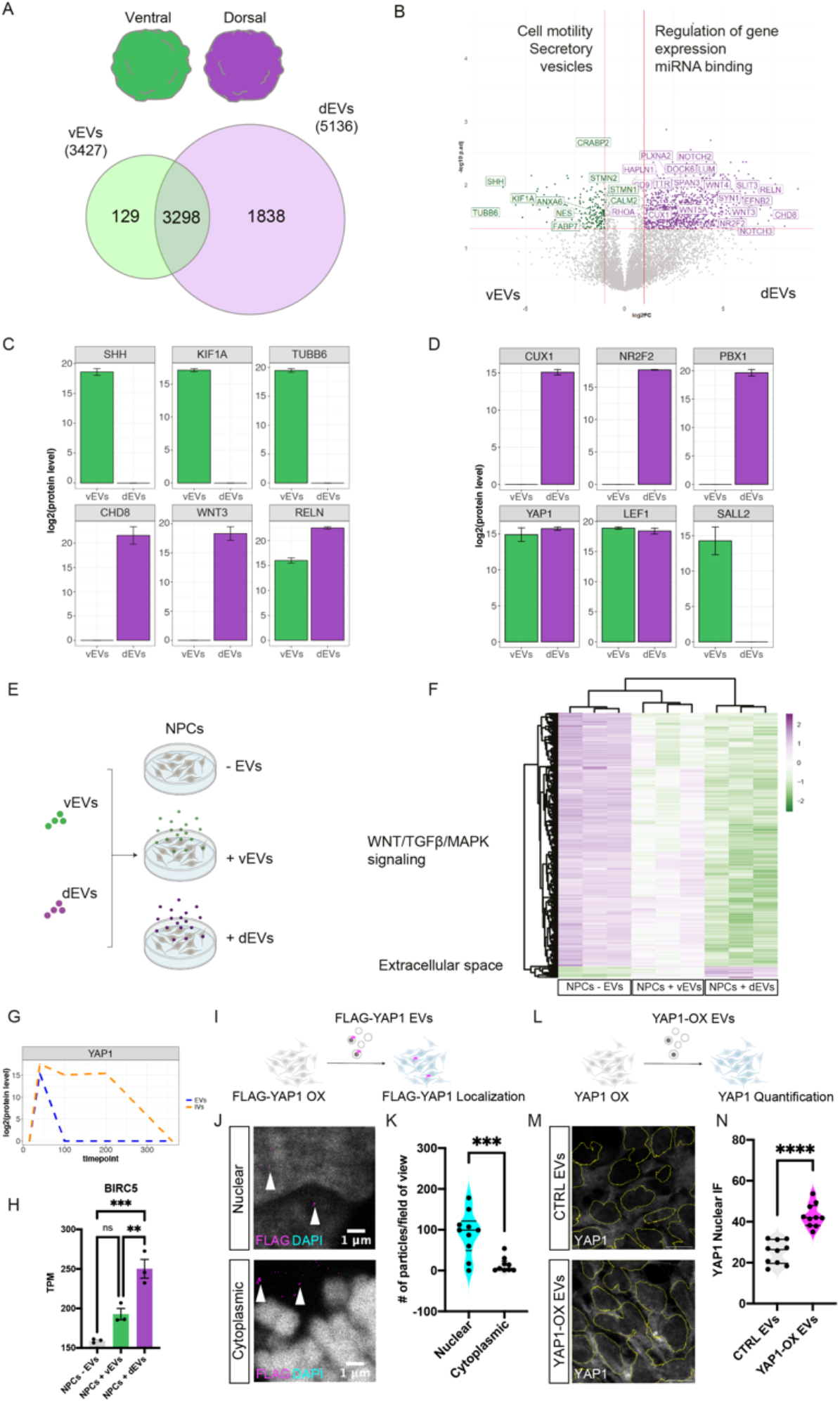
Brain-region-dependent signaling function of EVs. (A) Schematic of brain-region-specific COs. vCOs (ventral, green), dCOs (dorsal, purple) (top) and Venn diagram of vCO EVs (vEVs) and dCO EV (dEVs) proteins (bottom).(B) Volcano plot of protein level of vEVs and dEVs from 40days old COs, plotting the negative log10 q-values (FDR) of all proteins against their log2 fold change (dEVs vs vEVs). Significantly expressed proteins (q-value < 0.05) are labelled (vEVs, green and dEVs, purple). GO enrichments are shown. EVs were collected from the conditioned media of 15cm petri dishes containing 20-30 COs. (C) Bar plots showing the expression of brain-region-specific markers in vEVs and dEVs. Data are represented as mean ± SD of technical replicates. n= 2 CTRL lines and 2 EPM1 lines. (D) Bar plots showing the expression of transcriptions factors in vEVs and dEVs. Data are represented as mean ± SD of technical replicates. n= 2 CTRL lines and 2 EPM1 lines. (E) Schematic of acute treatment (12h) of NPCs with brain-region-specific CO EVs (vEVs, ventral, green; dEVs, dorsal, purple). (F) Heatmap showing differentially regulated genes in NPCs after treatment with vEVs and dEVs versus no EVs. n= 3 replicates per condition. (G) Temporal trajectory of YAP1 expression in EVs and IVs at different stages. (H) Bar plot showing the expression (TPM, Transcripts Per Million) of BIRC5 in NPCs after treatment with no EVs, vEVs, and dEVs. Data are represented as mean and ± SEM. n=3 (per condition). Statistical significance is based a one-way analysis of variance (ANOVA), **p<0.01, ***p<0.001. (I) Schematic of treatment with FLAG-YAP1 EVs derived from FLAG-YAP1 overexpressing NPCs for the localization of FLAG-YAP1 in receiving NPCs. (J) Immunostaining indicating uptake of FLAG-YAP1 EVs (magenta) by NPCs (Nuclear, top; Cytoplasmic, bottom). Arrow heads point to EV localization. DAPI, cyan. Scale bars: 1 μm. (K) Quantification of the number of particles detected in NPCs following treatment with FLAG-YAP1 EVs versus treatment with control EVs. Data are represented as mean and ± SEM. Every dot refers to a field of view, n= 10 (nuclear), n= 9 (cytoplasmic). Statistical significance is based a Student’s t-test, ***p<0.001. (L) Schematic of treatment with FLAG-YAP1 EVs derived from FLAG-YAP1 overexpressing NPCs for the quantification of YAP1 nuclear expression in receiving NPCs. (M) Immunostaining of YAP1 in NPCs treated with FLAG-YAP1 EVs with a yellow outline delimiting the cell nuclei. DAPI, cyan. Scale bars: 10 μm. (N) Quantification of nuclear YAP1 fluorescence intensity detected in NPCs following treatment with FLAG-YAP1 EVs versus treatment with control EVs. Data are represented as mean and ± SEM. Every dot refers to a field of view, n= 10 (per condition). Statistical significance is based a Student’s t-test, ****p<0.0001. Illustrations created using BioRender.

Amongst the unique proteins, typical patterning-related proteins were expressed either in dEVs (WNT3A) or vEVs (SHH) (Fig. 3C). An essential molecular motor (KIF1A) and other proteins associated to neurodevelopmental disorders (RELN) were specific or enriched in dEVs or vEVs (Fig. 3C). Single-cell RNA-Seq of dCOs and vCOs indicated a similar patterned expression of SHH and KIF1A compared to their EV expression (Fig. S4B-C). On the contrary, TUBB6 and CHD8 showed a broader RNA expression but a patterned EV protein load (Fig. 3C and S4B-C). Interestingly dEVs contained 84 transcription factors (TFs), including TFs fundamental during neurogenesis (examples in Fig. 3D) while vEVs only 50, of which 48 were shared with dEVs (examples in Fig. 3D). The levels of TFs loaded into EVs did not strictly correspond with their expression levels (Fig. S4B-C), suggesting a regulated secretion of TFs by specific cell types. To dissect if EVs, and particularly the TFs contained in EVs, have a functional role on cellular crosstalk, we investigated transcriptional changes on cells exposed to EVs. We performed RNA-seq analysis on NPCs acutely treated (12 hours) with EVs from dCOs and vCOs (Fig. 3E). NPC transcriptome was significantly altered upon EV treatment, particularly upon treatment with dEVs compared to vEVs (Fig. 3F).

To identify if some of the transcriptional changes observed in the NPCs that were exposed to EVs were caused by the TFs contained in vEVs and dEVs, we analyzed the expression of some of the TFs’ targets. BIRC5 (target of YAP1), ECT2 and RACGAP1 (targets of CUX1), HEY1 and HEY2 (targets of NR2F2) were significantly altered according to the TFs enrichment in the EVs (Fig. 3D, G-H and S4D-E). As YAP1 is known to be highly expressed (Sahu and Mondal, 2021) and it is secreted in NPCs at early stages of human development (Fig. 3G), we tested if we could track its journey, from secretion to uptake. To achieve this, donor NPCs were transfected with a FLAG-YAP1 plasmid, and receiving NPCs were treated with the EVs collected from their medium, containing FLAG-YAP1 (Fig. 3I-L). Immunostaining with FLAG and YAP1 antibodies performed in the cells exposed to the FLAG-YAP1-EVs revealed the presence of FLAG in the cytoplasm and nucleus of the receiving NPCs and increased YAP1 intensity compared to treatment with control EVs (Fig. 3 J-N).

Together these data suggest that EVs play a role during development as they contain regulators such as TFs, that can be translocated from cell to cell and lead to a clear transcriptional change in receiving cells.

### Impact of EVs on neurodevelopmental disorders

Recent evidence suggests that neurodevelopmental disorders are associated to changes in the extracellular environment (Amin and Borrell, 2020; Mazurskyy and Howitt, 2021). 23% of proteins coded by genes associated to some neurodevelopmental disorders, particularly Cortical Malformations (CMs), Epilepsy, Autism Spectrum Disorders (ASD) and Schizophrenia (SCZ) were found in vesicles (Fig. 4A and S4F). Comparison of neurodevelopmental disorders genes and EV-proteins in different cell types clearly suggested a neuron/late-enrichment in disease secreted proteins (Fig. S4G-I).

**Fig. 4.**
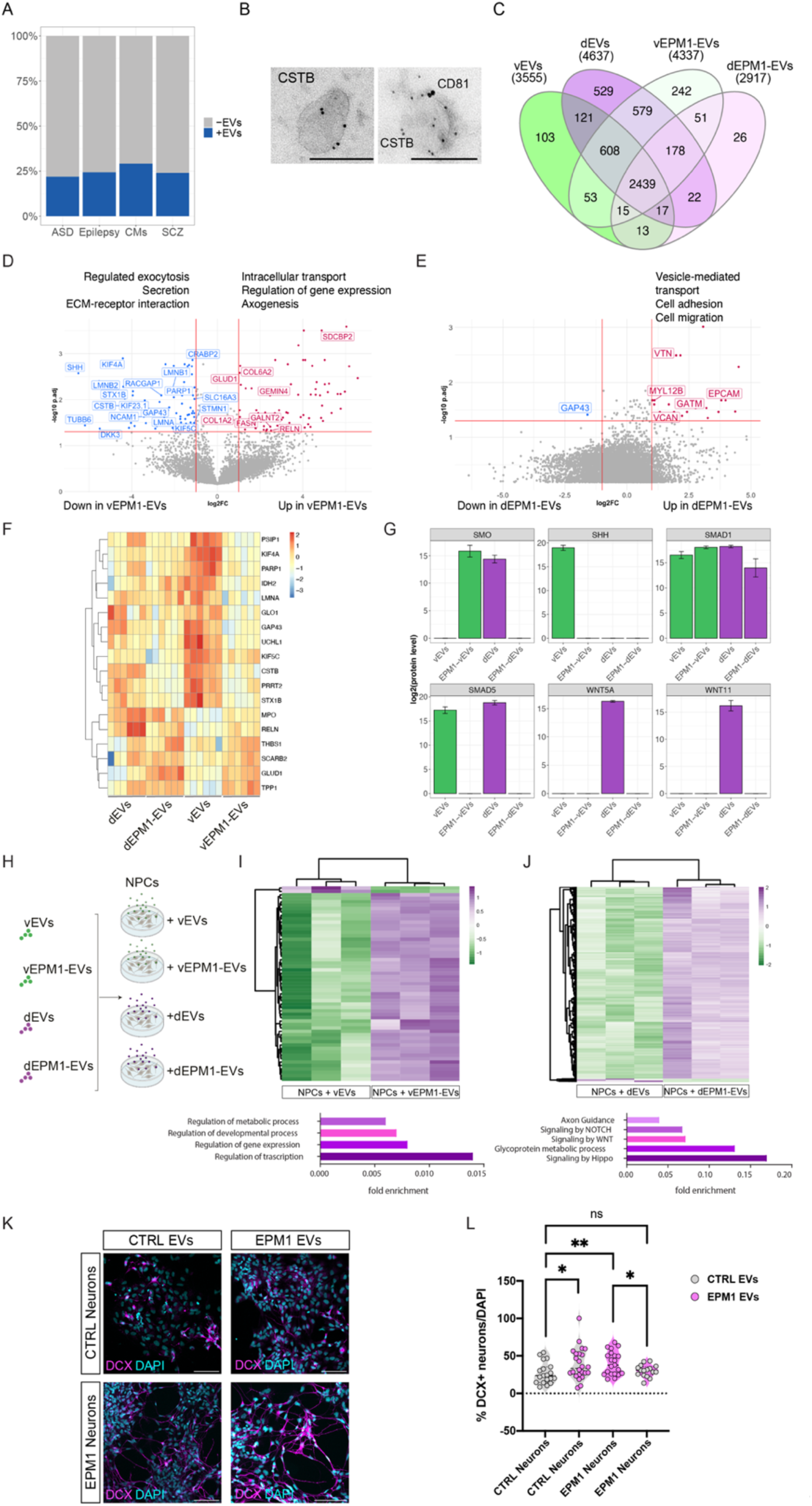
Impact of EVs on neurodevelopmental disorders. (A) Bar plot indicating the proportion of proteins found in CO EVs associated with some neurodevelopmental disorders (DisGeNET). (B) Immuno-electron micrographs of CD81 (large dots) and CSTB (small dots) in EVs collected from COs. Scale bar: 100nm. (C) Venn diagram of vEVs, dEVs, vEPM1-EVs and dEPM1-EVs proteins from 40day old COs. (D) Volcano plot of protein level of vEVs and vEPM1-EVs, plotting the negative log10 q-values (FDR) of all proteins against their log2 fold change (vEPM1-EVs vs vEVs). Significantly expressed proteins (q-value < 0.05) are labelled. GO enrichments are shown. EVs were collected from the conditioned media of 15cm petri dishes containing 20-30 COs. EV proteomic analysis was performed on EPM1 patient-derived COs from two different patients (pooled together for this analysis). (E) Volcano plot of protein level of dEVs and dEPM1-dEVs, plotting the negative log10 q-values (FDR) of all proteins against their log2 fold change (dEPM1-dEVs vs dEVs EVs). Significantly expressed proteins (q-value < 0.05) are labelled. GO enrichments are shown. EVs were collected from the conditioned media of 15cm petri dishes containing 20-30 COs. EV proteomic analysis was performed on EPM1 patient-derived COs from two different patients (pooled together for this analysis). (F) Heatmap showing the 289 vEV, dEV, vEPM1-EV and dEPM1-EV proteins associated with epilepsy. (G) Bar plots showing the protein levels of brain patterning molecules in vEVs, vEPM1-EVs, dEVs, and dEPM1-EVs. Data are represented as mean ± SD of 3 technical replicates. EVs were collected from the conditioned media of 15cm petri dishes containing 20-30 COs. EV proteomic analysis was performed on EPM1 patient-derived COs from two different patients (pooled together for this analysis). (H) Schematic of acute treatment (12h) of NPCs with brain-region-specific CO and EPM1-EVs (vEVs and vEPM1-EVs, green; dEVs and dEPM1-EVs, purple). Illustration created using BioRender. (I) Heatmap showing z-scores of the expression levels of the 57 differentially regulated genes in NPCs after acute treatment with vEVs and vEPM1-EVs and with (J) dEVs and dEPM1-EVs (1314 differentially regulated genes). P.adj < 0.05, fold change > 2; n = 3 for each condition. (K) Immunostaining of DCX+ CTRL and EPM1 neurons after treatment with CTRL or EPM1 EVs. Scale bar: 100 µm. (L) Quantification of the percentage of DCX+ neurons/DAPI (CTRL vs EPM1) after treatment with CTRL or EPM1 EVs. Data are represented as mean and ± SEM. Every dot refers to a field of view, n= 16-24 per condition. Statistical significance is based on multiple t-tests *p<0.05; **p<0.01; ns, not significant.

Progressive Myoclonus Epilepsy Type I (EPM1) is a rare type of epilepsy that arises from mutations in *CSTB*, a gene which codes for Cystatin B, a ubiquitous protease inhibitor (Lalioti et al., 1997). Some of the proteins belonging to the cystatin family have already been shown to be secreted (Di Matteo et al., 2020; Ochieng and Chaudhuri, 2010); however, the cell-type specificity of their released in EVs has not been studied. Thus, we analyzed if cystatin proteins are contained in EVs from different 2D cell cultures. Interestingly, only CSTA and CSTB were detected in EVs of mutually exclusive cell types (Fig. S4J). Proteins of the cystatin family are expressed in EVs with varying temporal dynamics (Fig. S4K). CSTA and CSTB again show mutual-exclusive expression, while the expression of CSTB and CST3 in EVs overlaps at later stages of development. Interestingly, CST3 overexpression has a beneficial effect in mice lacking CSTB (Kaur et al., 2010). Meanwhile, we observed that both vEVs and dEVs contain CSTA, CSTB and CST3 proteins at similar levels (Fig. S4L). To investigate if CSTB contributes to signaling mechanisms mediated by EVs, we first performed immuno-EM to verify the presence of CSTB in EVs. We confirmed CSTB expression in a variety of EVs, more specifically CD81+ and CD81-vesicles (Fig. 4B). We performed EV proteomic analysis from dorsal and ventral EPM1 patient-derived COs (EPM1-COs) from two different patients (pooled together for this analysis) as previously described (Di Matteo et al., 2020)(Fig. 4C-E, S5A) compared to control COs from two control iPSC lines. Strikingly, we found a strong decrease of proteins in EPM1-dEVs compared to dEVs (37,1%) and an increase in EPM1-vEVs compared to vEVs (22%, Fig. 4C). By clustering the EPM1-EV DE proteins with the protein content of pEVs, nEVs and aEVs, we found an enrichment for proteins transported in EVs secreted by NPCs (Fig. S5B). Intriguingly, EVs from EPM1-dEVs critically lack multiple of the TFs contained in dEVs and vEVs, suggesting that EPM1-dEVs are depleted from essential signals for cell-to-cell communication. Following DE expression analysis with DESeq2, GAP43, a growth-cone enriched protein in maturing neurons, was significantly decreased in EPM1-dEVs. Moreover, extracellular matrix-receptor interaction, secretion and regulation of gene expression were dysregulated in EPM1-vEVs, while vesicle-mediated transport and cell adhesion were dysregulated in EPM1-dEVs, indicating that CSTB is critical for appropriate cellular environment and gene expression (Fig. 4D-E). One of the most intriguing results from the proteomics analysis presented is that 18 proteins altered in EPM1-EVs, are coded by genes whose mutations are associated to epilepsy (Fig. 4F). Epilepsy is often caused by an imbalance between excitatory and inhibitory neurons, as is specifically in the case of EPM1. We therefore hypothesized that the proteins loaded into EVs and transported locally could influence the specification and identity of excitatory or inhibitory neurons. Strikingly, we identified a higher number of significantly differentiated proteins amongst ventral EVs (Fig. 4D). Amongst the differentially expressed proteins in vEVs, we found essential components of the SHH, SMAD and WNT pathways (SMO, SHH, SMAD1, SMAD5, WNT5A and WNT11), required for progenitors and neuron specification (Takahashi and Liu, 2006) (Fig. 4G). To investigate if these major modifications in EV content directly triggered changes at the transcriptional level in receiving cells, we collected EVs from EPM1-COs and treated control NPCs acutely (12 hours, schematic in Fig. 4H) with dEVs, vEVs, EPM1-dEVs and EPM1-vEVs (Fig. 5H). We then performed transcriptomic analysis of the recipient NPCs (Fig. 4I-J and S5C). Upon treatment with EPM1-EVs, both dorsal and ventral, most of the NPC altered genes were upregulated compared to treatment with control EVs. As previously described, dEVs mediated most of the transcriptional changes, including genes associated with the Hippo pathway. Only 2 genes (KRT8 and KRT19) were downregulated in NPCs treated with EPM1-vEVs and 22 were overexpressed upon EPM1-dEVs treatment. These data further suggest that EPM1-EVs lack critical signals for cell-to-cell communication. Transcriptome changes of NPCs treated specifically with EPM1-vEVs included 57 differentially expressed genes of which 17 relate to the SHH pathway. To assess the functional role of EVs in EPM1 epilepsy and because of the altered SHH pathway in both donor EVs and receiving NPCs, we generated EPM1-vCOs and analyzed the presence of ventral and dorsal markers at 30d in culture. EPM1-vCOs showed a strong decrease in NKX2.1+ ventral ventricles and a significantly increased number of TBR1+ dorsal ventricles (Fig. S5D-G). To investigate if these changes were triggered by a non-cell autonomous mechanism, we generated hybrid vCOs containing GFP-labeled CTRL cells and non-labeled EPM1 cells. CTRL (GFP+), EPM1 (GFP-) and hybrid (GFP+/-) ventricles were then analyzed with the ventral progenitor markers NKX2.1 and MEIS2 (Fig. S5H-K). Progenitors contained in the germinal zone of the hybrid ventricles showed an identity closer to EPM1 ventricles, suggesting that critical levels of SHH and other proteins contained in EVs are essential for proper progenitor patterning in the developing brain. As most of the interneurons residing in the adult brain originate from ventral progenitors, these fate changes in early stages of development may be crucial for the proper establishment of a correct excitation/inhibition balance. Additionally, as it was previously shown that EPM1 neurons differentiate prematurely (Di Matteo 2020), we treated young control neurons with EPM1-EVs and young EPM1 neurons with control EVs for two weeks. The EV exchange showed a pathological function of EPM1-EVs as control neurons differentiate faster upon treatment with EPM1-EVs (Fig. 4K-L). These results suggest that EVs contain cargoes for progenitor and neuronal specification and differentiation.

**Fig. 5.**
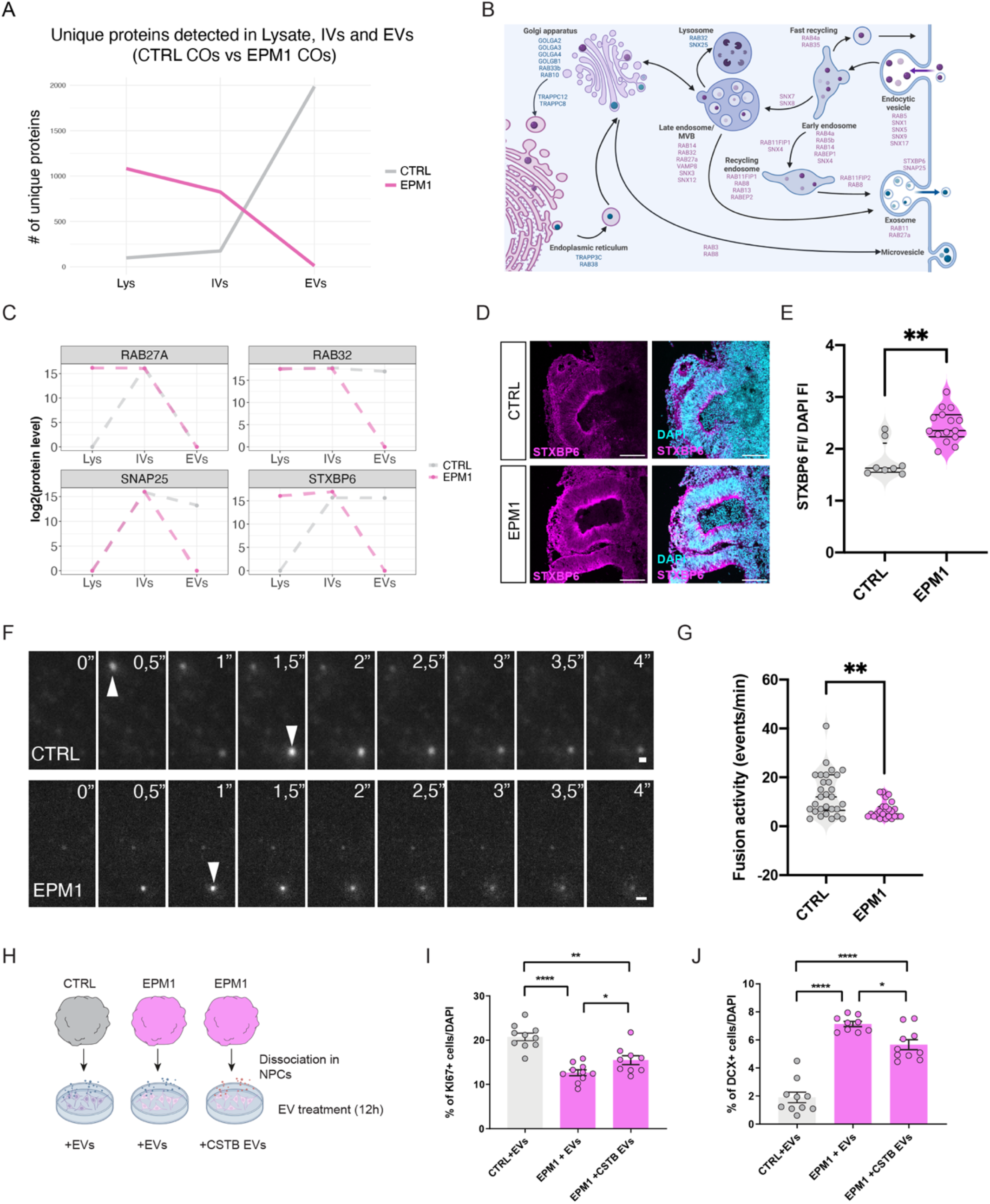
Extrinsic role of vEPM1-EVs on cell fate. (A) Line graph showing the unique number of proteins detected in Lysates (Lys), IVs and EVs from CTRL and EPM1 COs. EVs were collected from the conditioned media of 15cm petri dishes containing 20-30 COs. EV proteomic analysis was performed on EPM1 patient-derived COs from two different patients (pooled together for this analysis). Total cell lysates and IVs were obtained from a pool of 5-7 COs. (B) Schematic of the dysregulated proteins in EPM1-EVs that are involved in EV trafficking. (C) Line graphs of the protein level of RAB27A, RAB35, RAB31 and STXBP6 in the cell lysate, IVs and EVs. EVs were collected from the conditioned media of 15cm petri dishes containing 20-30 COs. EV proteomic analysis was performed on EPM1 patient-derived COs from two different patients (pooled together for this analysis). Total cell lysates and IVs were obtained from a pool of 5-7 COs. (D) Immunostaining of STXBP6 (magenta) in CTRL COs and EPM1-vCOs at 30d. DAPI (cyan). Scale bar: 100 µm.(E) Quantification of STXBP6 fluorescence intensity/ DAPI in CTRL COs and EPM1-vCOs at 30d. Data are represented as mean and ± SEM. Every dot refers to a CO ventricle, n= 8 (CTRL), n= 16 (EPM1), 6-8 COs were analysed per condition. Statistical significance is based on a Mann-Whitney U test *p<0.01. (F) Time lapse images of MVB-plasma membrane fusion events (arrow heads) in CTRL and EPM1 neurons. Scale bar: 1 µm. (G) Quantification of the fusion activity (events/min) detected in CTRL and EPM1 neurons. Data are represented as mean and ± SEM. Every dot refers to a single cell analyzed, n= 29 (CTRL), n= 26 (EPM1). Statistical significance is based on a Mann-Whitney U test *p<0.01. (H) Experimental setup of NPCs (derived from control and EPM1 COs) treated for 7 days with control EVs and +CSTB EVs. (I) Quantification of KI67+ NPCs (derived from control and EPM1 COs) treated as shown in (J). Data are represented as mean and ± SEM. Every dot refers to a field of view, n= 9-10 (per condition). Statistical significance was based on the Mann-Whitney U test *p<0.05, **p<0.01, ****p<0.0001. (K) Quantification of DCX+ NPCs (derived from control and EPM1 COs) treated as shown in (J). Data are represented as mean and ± SEM. Every dot refers to a field of view, n= 9-10 (per condition). Statistical significance was based on the Mann-Whitney U test *p<0.05, **p<0.01, ****p<0.0001. Illustration created using BioRender.

### Intracellular/extracellular trafficking is altered in EPM1

CSTB is a ubiquitous protease inhibitor that reduces the activity of enzymes of the cathepsin family. The most characterized function of cathepsins is the regulation of protein breakdown in lysosomes (Yadati et al., 2020). Lysosomes can fuse with endosomes, phagosomes, autophagosomes, and breakdown both endogenous and exogenous cargo consisting of various biomolecules such as lipids, proteins, polysaccharides, and certain pathogens (Yadati, Cells 2020). The known functions of cathepsins, together with our finding that the cargo loaded in EPM1-EVs is altered, led us to the hypothesis that CSTB is involved in protein trafficking. We therefore examined the unique protein content in different cellular compartments, including vesicles inside (IVs) and outside (EVs) of the cells. Strikingly, a comparison of proteins in different compartments highlighted a number of unique proteins present only in control or EPM1 conditions. While the total number of unique proteins found in cells and IVs was higher in EPM1 organoids, the number of proteins in EVs was strongly reduced (Fig. 5A), suggesting alterations in EV biogenesis and secretion in patient organoids. Amongst these unique proteins we found an interesting enrichment for proteins involved in trafficking, including several proteins of the RAB, SYN, STX, TRAPP, GOLG and SNARE families (Fig. 5B).

Interestingly, RAB27A, RAB32, SNAP25 and STXBP6 are important players for the regulation of EV secretion (Kondratiuk et al., 2020; Nassari et al., 2020) (Fig. 5C). A possible role of CSTB in vesicular trafficking and autophagy was proposed in astrocytes isolated from CSTB knock-out mice where levels of CD63 were increased compared to control astrocytes (Polajnar et al., 2014). Interestingly, one of the few upregulated proteins in EPM1 organoids was the small SNARE protein STXBP6 that acts as a vertebrate-specific competitor of synaptobrevin-2, key player in membrane fusion during exocytosis (Kondratiuk et al., 2020) (Fig. 5D-E and S5L). To demonstrate our hypothesis that EV secretion is altered in EPM1 conditions, we monitored the membrane fusion activity of control and EPM1 neurons by live imaging using TIRF microscopy after overexpressing CD63-pHluorin, a tetraspanin-based pH-sensitive optical reporter that detects multivesicular body -plasma membrane (MVB-PM) fusion (Verweij et al., 2018) (Fig. 5F-G). EPM1 neurons showed a decreased number of events/minute, indicating an altered EV secretion in pathological conditions (Fig. 5G). Taken together EPM1 patients showed an altered composition, function and dynamic of secretion of EVs.

These results in principle suggest that manipulation of the content of EVs could be a new avenue for developing strategies in order to rescue some of the EPM1 phenotypes. To test this idea, we overexpressed CSTB in donor SH-SY5Y cells and collected EVs (CSTB-EVs) from the conditioned media.

Upon treatment with CSTB-EVs, EPM1 cells increased their proliferation (KI67+ cells) and decreased their differentiation (DCX+ cells), suggesting that the presence of CSTB was sufficient to load the correct cargoes into EVs that triggered these changes and promoted a partial rescue (Fig. 5H-J and S5M). Intriguingly, 40% of the CSTB-EV enriched protein content consists of downregulated proteins in EPM1-EVs derived from COs, suggesting an important role of CSTB in the correct biogenesis (protein loading) and function of EVs.

## Conclusion

In this study, we characterized the developmental-, regional-, cell-type-specific protein composition of EVs in human COs, highlighting the specific function of EVs in physiological and pathological conditions.

We have presented the first evidence that providing a 3D environment during development is critical for building heterogeneous EVs, therefore emphasizing the contribution of tissue complexity to the landscape of EVs in the extracellular space. Moreover, our data indicate that different cell types use specific mechanisms to receive signals from EVs. These differences in composition and uptake may be responsible for a unique cell-specific crosstalk during brain development. Additionally, we present changes in EV composition and function in a CO model of epilepsy, highlighting a potential mechanism for alterations in the extracellular environment that may lead to changes in cell fate.

This knowledge provides novel insight into cell non-autonomous mechanisms involved in human brain development that could be disturbed in neurodevelopmental disorders. Taken together, our results could lead to advances in new therapeutic strategies for patients exhibiting epilepsy.

## Materials and Methods

### IPSCs culture

iPSCs were previously reprogrammed from 2 control lines of fibroblasts (Klaus et al., 2019) and PBMCs origin (Di Matteo et al., 2020) and from 2 EPM1 patient lines (Di Matteo et al., 2020). iPSCs were cultured on Matrigel (Corning) coated plates (Thermo Fisher, Waltham, MA, USA) in mTesR1 basic medium supplemented with 1x mTesR1 supplement (Stem Cell Technologies, Vancouver, Canada) at 37°C, 5% CO2 and ambient oxygen level. Passaging was done using accutase (Stem Cell Technologies) treatment.

### Generation of labelled iPSC lines

The RFP-labeled iPSC lines were generated using the piggyBac transposase (1 ug) and PB-RFP (1ug) nucleofection (F and J, 2012). Single cells of iPSCs were transfected with the Amaxa Nucleofector 2b (program B-016). RFP positive colonies were picked and cultured on Matrigel (Corning/VWR International, 354234) coated plates in mTeSR1 basic medium (Stem Cell Technologies, 85850) supplemented with 1× mTeSR1 supplement (Stem Cell Technologies, 85850) at 37°C and 5% CO2.

### Generation of human cerebral organoids

Reprogrammed iPSCs were used to generate human cerebral organoids (hCOs) as previously described (Lancaster and Knoblich, 2014; Lancaster et al., 2013). Organoids were kept in 10-cm dishes on a shaker at 37°C, 5% CO2 and ambient oxygen level with medium changes every 3–4 days.

### Generation of patterned human organoids

Patterned human organoids were generated according to (Bagley et al., 2017). Embryoid bodies (EBs) generated from iPSCs were patterned to have ventral and dorsal identity. During the neuronal induction step, EBs were treated individually with SAG (1:10,000) (Millipore, 566660) + IWP-2 (1:2,000) (Sigma-Aldrich, I0536) for inducing ventral identity, s with cyclopamine A (1:500) (Calbiochem, 239803) for inducing dorsal identity. After this point, the generation of organoids followed methods according to (Lancaster and Knoblich, 2014).

### Generation of hybrid human organoids

Ventral hybrid COs were generated according to Bagley et al, 2017. iPSCs from GFP-labeled iPSC control line and from EPM1 iPSCs, were dissociated into single cells using Accutase (Sigma-Aldrich, A6964), mixed together in a ratio of 30% GFP-control cells : 70% EPM1 cells and transferred in a total of approximately 9000 cells to one well of an ultra-low-attachment 96-well plate (Corning). The protocols continued as described in “Generation of patterned Human organoids”.

### NPCs, neurons and astrocytes cultures

Neural progenitor cells (NPCs) were generated and cultured by following Basic Protocol 1 as previously described (Boyer et al., 2012), with the exception that FGF8 and SHH were replaced by FGF2 (Peprotech, 100-18b-50) in the neural progenitor medium (NPM). NPCs were generated from two control iPSC lines, one RFP-labelled line and one unlabelled line (see “IPSCs culture” and “Generation of labelled iPSC lines”), which generated a ratio of 60% neurons and 40% astrocytes in accordance with this protocol, providing electrophysiologically mature neurons in a more physiological environment. Neural differentiation was conducted as previously described (Gunhanlar et al., 2017). Astrocytes were isolated from 8-month-old organoids as follows: Organoids were transferred to a 15 ml falcon tube and washed 1 time with 1xPBS. For dissociation, they were placed in Accutase® solution (A6964, Sigma Aldrich, St. Louis, MO, USA) pipetted up and down 5-10 times with a P1000 tip, and then placed in the incubator for 10 mins at 37°C, followed by 5 times pipetting for a second time. The dissociated cells were then centrifuged at 300 x g for 3 mins and resuspended in *NDM+A media* (DMEME/F12+Glutamax and Neurobasal™ medium in a ratio 1:1 supplemented with 1:100 N2™-supplement (100X), 1:100 B-27™ supplement (50X), 0.5% of MEM Non-Essential Amino Acids Solution (100X), 0.5% GlutaMAX™ Supplement, 50 uM of 2-mercaptoethanol (50 mM), antibiotic antimycotic Solution (100×) and Insulin 2.5 ug/ml) for 24 hours. The next day, the cells were transferred to Matrigel® Basement Membrane Matrix LDEV-free (Corning®, 354234) coated plates. One day later the media was changed to Astrocyte media (89% DMEM/F12+Glutamax, 10% fetal bovine serum, 1% Antibiotic-Antimycotic). The astrocytes obtained were characterized by immunostaining and were positive for astrocytic markers such as SOX9, s100B, NFIA, and negative for neuronal markers MAP2 and NeuN. All the cells were kept in an incubator at 37°C, 5% CO2 and ambient oxygen level with medium changes every 2–3 days.

### Exchange of EVs from conditioned media from CTRL and EPM1 neurons

CTRL and EPM1 NPCs were differentiated to neurons for 2 weeks (see “NPCs, neurons and astrocytes cultures”). During these 2 weeks, receiving neurons were treated with EVs collected from conditioned media collected from donor cultures of CTRL and EPM1 neurons. EVs were collected by the following steps: conditioned media centrifugation at 300 g for 15 mins, supernatant centrifugation at 2000g for 10 mins, supernatant centrifugation at 100,000 g for 90 mins. CTRL and EPM1 EVs were added to receiving neurons, either CTRL or EPM1, every third day until the day of the analysis. At 2 weeks, receiving neurons were collected for immunohistochemistry analysis and imaged using a confocal microscope (see Immunohistochemistry and Imaging). The images were then analysed using ImageJ (Schneider et al., 2012).

### FLAG-YAP1 overexpression and EV exchange

NPCs were transfected with a FLAG-YAP1 plasmid (Addgene #27371, (Cappello et al., 2013)) and a control plasmid (pEGFP-C1 plasmid) using the Lipofectamine™ 3000 Transfection Reagent (ThermoFisher Scientific, USA) as instructed in the protocol. 72h following the transfection, the conditioned media from the NPCs was collected for EV isolation. The collected EVs were then added to a new set of NPCs, and the cells were prepared for immunofluorescence after 18 hrs of the treatment and imaged using confocal microscopy (see Immunohistochemistry and Imaging). The images were then analysed using ImageJ (Schneider et al., 2012).

### Transfection of SH-SY5Y cell line

SH-SY5Y cells were cultured in DMEM/F12GlutaMAX medium supplemented with 10% FBS and 1% antibiotics at 37°C, 5% CO2 and ambient oxygen level. Passaging was done using trypsin/EDTA (Sigma) treatment. For EVs collection, one day before the start of the experiment (day -1) cells were cultured in media with exosome depleted FBS (Gibco, 15624559). The day 0, 2 million cells (80% confluency) were cultured in 15 cm plates in DMEM/F12GlutaMAX medium supplemented with 10% FBS (exosomes depleted) without antibiotics and overexpression of GFP and GFP-CSTB *(18)*, was performed via electroporated with 4 ugr of pEGFP-C1 expression construct using the Amaxa nucleofector at the program G004. The following day (day 1) antibiotics were added to the medium. On Day 2 the conditioned medium was collected for EVs purification.

### EVs and IVs collection and analysis

In accordance with (Théry et al., 2006) EVs were collected from conditioned media from COs and 2D cultured cells by the following steps: centrifugation at 300g for 10 mins, supernatant centrifugation at 2000g for 10 mins at 4 °C, supernatant centrifugation at 10.000g for 30 mins at 4 °C, supernatant centrifugation at 100.000g for 90 mins at 4 °C in a fixed-angle rotor (TH865, Thermo Fisher Scientific), followed by pellet wash with 1x PBS and centrifugation at 100.000g for 90 mins at 4 °C. Alternatively, miRCURY Exosome Cell/Urine/CSF Kit (Qiagen, 76743) was used to isolate EVs from conditioned medium according to the manufacturer instructions. For NPCs, EVs were collected from three independent cultures of control NPCs. Neuronal EVs were collected from three independent neuronal differentiation cultures. Similarly, astrocyte EVs were collected from three independent cultures of astrocytes. For COs, EVs were collected from conditioned media of 20-30 different COs in culture.

IVs were isolated by subcellular fractionation. Briefly, a pool of 5-7 COs were homogenized and upon removal of nuclei, cell debris and mitochondrial fraction as previously reported (Ferrara et al., 2009), the supernatatant was ultracentrifuged at 100.000 g for 30 min to obtain the cellular fraction (Ivs).

For the nanoparticle tracking analysis (NTA), fresh, unfrozen extracellular vesicle suspensions were diluted in PBS and analysed using a Particle Metrix ZetaView® PMX110-Z Nanoparticle Tracking Analyzer (Particle Metrix GmbH, Inning am Ammersee, Germany) equipped with a 520 nm laser. For each measurement, samples were introduced manually, the temperature was set to 24 °C, and two cycles were performed by scanning at 11 discrete positions in the cell channel and capturing 60 frames per position (video setting: high). The following recommended parameters were used for the measurement:

Sensitivity: 80.0

Shutter: 70

Frame rate: 30

Minimum 20

Brightness:

Minimum Area: 5

Maximum Area: 1000

Maximum Brightness: 255

Brightness:

Tracking Radius2: 100

Minimum 15

Tracelength:

nm/class: 5

Classes/decade: 64

After capture, the videos were analysed for particle size and concentration using the ZetaView Software 8.05.12 SP1.

For immune-electron microscopy, aliquots of extracellular vesicle suspensions were anayzed by Dr Ilkka Miinalainen at Biocenter Oulu / EM laboratory, Finland (Deun et al., 2020). Vesicles were deposited on Formvar carbon coated, glow-discharged grids and incubated in a blocking serum containing 1% BSA in PBS. CD81, LGALS3, CSTB primary antibodies and secondary gold conjugates (Zymed, San Francisco, CA, USA) were diluted in 1% BSA in PBS. The blocking efficiency was controlled by performing the labelling procedure in the absence of primary antibody.

### Dissociation of ventral and dorsal control organoids for 2D cultures

2 months old ventral and dorsal hCOs generated from control and EPM1 iPSCs, were dissociated as previously described (Di Matteo et al., 2020), with some modifications. Briefly hCOs were dissociated to single cells using Accutase (Sigma-Aldrich, A6964). Single cells were then plated onto Poly-L-ornithine (10 μg/ml) (Sigma-Aldrich, P4957)/Laminin (10 μg/ml) (Sigma-Aldrich, L2020)-coated coverslips with some modifications in wells of 24-well plates (Corning). Cells coming from a pool of two to three organoids were plated in 12 wells of 24-well plates with neural progenitor cells medium (NPC medium) (Gunhanlar et al., 2017). After 7 days in culture, cells were treated with GFP or GFP-CSTB EVs for 7 days. Freshly collected EVs from culture media of transfected SH-SY5Y cells were added directly to the media during the media changing (3 times per week). After 14 days in culture, cells were fixed with 4% PFA, and then, immunostainings were performed using KI67 to evaluate the number of proliferating cells and with doublecortin (DCX) to evaluate the number of young neurons derived from the organoids.

### EV uptake

NPCs, astrocytes and neurons were cultured in 24-well plates. 10-12 ml of conditioned media from astrocytes and neurons were treated with 1 ul of 10 mM DiI (1,1’-Dioctadecyl-3,3,3’,3’-Tetramethylindocarbocyanine Perchlorate) for 15 mins in the dark before the final washing step in ultracentrifugation. The media of the recipient cells was changed just prior to the addition of the labelled EVs. The cells were fixed 18 hours after EV treatment with 4% paraformaldehyde for 20 mins at room temperature.

### Immunohistochemistry

Cells dissociated from 2 months old COs were fixed using 4% PFA for 10 min and permeabilized with 0.3% Triton for 5 min. After fixation and permeabilization, cells were blocked with 0.1% Tween, 10% Normal Goat Serum (Biozol, VEC-S-1000). Primary and secondary antibodies were diluted in blocking solution. DCX, diluition 1:2000; Ki67 dilution 1:500. Nuclei were visualized using 0.5 mg/ml 4,6-diamidino-2-phenylindole (DAPI) (Sigma-Aldrich, D9542). Stained cells were analyzed using a Leica laser-scanning microscope.

NPCs, neurons, astrocytes and SH-SY5Y cells were fixed with 4% paraformaldehyde for 20 mins at room temperature, followed by three times 5 min washing with 1xPBS. Next, cells were blocked against unspecific binding and permeabilized in blocking buffer (10% normal goat serum, 0,02% Triton-X in 1xPBS) for 1 hour. Primary antibodies diluted in blocking buffer were then added to the cells in the dilutions specified below and incubated overnight. On the second day, cells were washed five times for 5 mins each in PBS with 0,1% Tween (PBS-T), and then incubated for 2 hours in secondary antibodies raised against the host animal of the primary antibody. Secondary antibodies were diluted in blocking buffer and the dilutions used are listed below (Table 1). DAPI (4′,6-diamidino-2-phenylindole) was added as a nuclear counterstaining. Finally, cells were washed three times with PBS-T and mounted on object slides with Fluoromount-G (ThermoFisher Scientific, 00-4958-02).

**Table 1.**
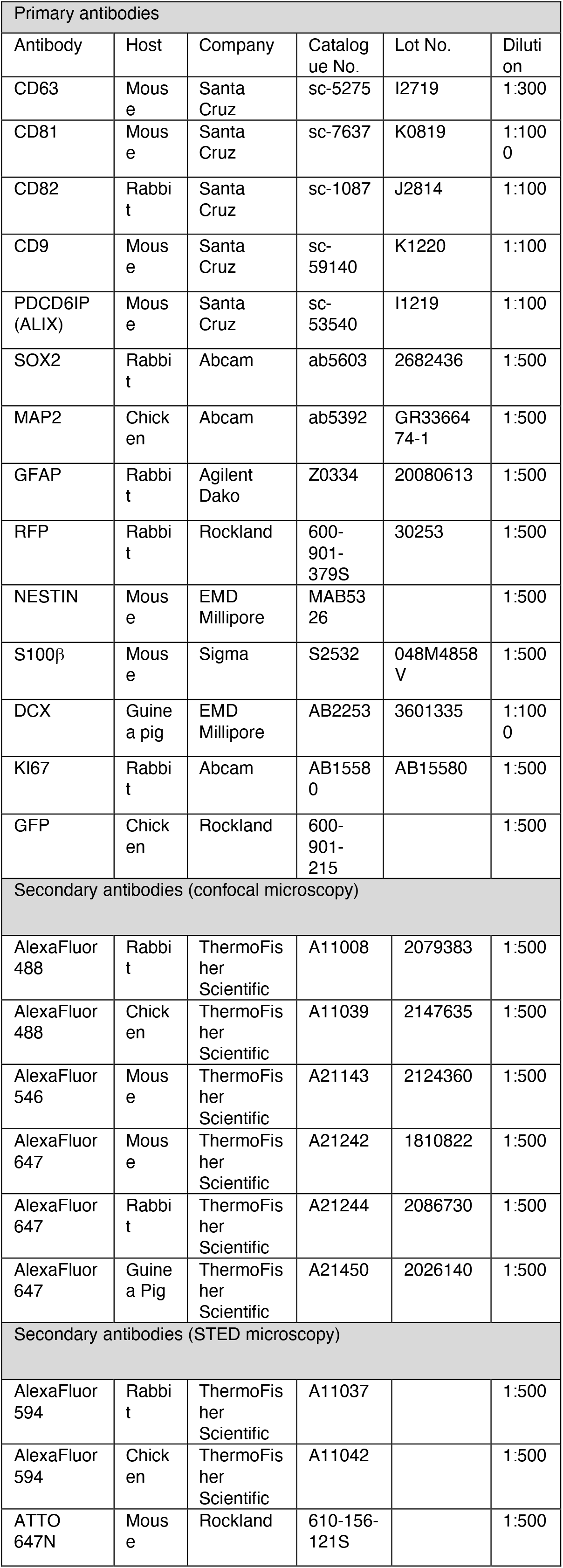
Immunostaining antibodies

| Primary antibodies |  |  |  |  |  |
| --- | --- | --- | --- | --- | --- |
| Antibody | Host | Company | Catalogue No. | Lot No. | Dilution |
| CD63 | Mouse | Santa Cruz | sc-5275 | I2719 | 1:300 |
| CD81 | Mouse | Santa Cruz | sc-7637 | K0819 | 1:1000 |
| CD82 | Rabbit | Santa Cruz | sc-1087 | J2814 | 1:100 |
| CD9 | Mouse | Santa Cruz | sc-59140 | K1220 | 1:100 |
| PDCD6IP (ALIX) | Mouse | Santa Cruz | sc-53540 | I1219 | 1:100 |
| SOX2 | Rabbit | Abcam | ab5603 | 2682436 | 1:500 |
| MAP2 | Chicken | Abcam | ab5392 | GR3366474-1 | 1:500 |
| GFAP | Rabbit | Agilent Dako | Z0334 | 20080613 | 1:500 |
| RFP | Rabbit | Rockland | 600-901-379S | 30253 | 1:500 |
| NESTIN | Mouse | EMD Millipore | MAB5326 |  | 1:500 |
| S100β | Mouse | Sigma | S2532 | 048M4858V | 1:500 |
| DCX | Guinea pig | EMD Millipore | AB2253 | 3601335 | 1:1000 |
| KI67 | Rabbit | Abcam | AB15580 | AB15580 | 1:500 |
| GFP | Chicken | Rockland | 600-901-215 |  | 1:500 |
| Secondary antibodies (confocal microscopy) |  |  |  |  |  |
| AlexaFluor 488 | Rabbit | ThermoFisher Scientific | A11008 | 2079383 | 1:500 |
| AlexaFluor 488 | Chicken | ThermoFisher Scientific | A11039 | 2147635 | 1:500 |
| AlexaFluor 546 | Mouse | ThermoFisher Scientific | A21143 | 2124360 | 1:500 |
| AlexaFluor 647 | Mouse | ThermoFisher Scientific | A21242 | 1810822 | 1:500 |
| AlexaFluor 647 | Rabbit | ThermoFisher Scientific | A21244 | 2086730 | 1:500 |
| AlexaFluor 647 | Guinea Pig | ThermoFisher Scientific | A21450 | 2026140 | 1:500 |
| Secondary antibodies (STED microscopy) |  |  |  |  |  |
| AlexaFluor 594 | Rabbit | ThermoFisher Scientific | A11037 |  | 1:500 |
| AlexaFluor 594 | Chicken | ThermoFisher Scientific | A11042 |  | 1:500 |
| ATTO 647N | Mouse | Rockland | 610-156-121S |  | 1:500 |

### Imaging

Immunostainings were imaged with confocal microscopy or Stimulated Emission Depletion (STED) microscopy. Confocal stack images were obtained using a Leica SP8 confocal microscope based on a DMi8 stand (Leica Microsystems, Wetzlar, Germany), equipped with 20x/0.75 (oil), 40x/1.10 (water), and 63x/1.30 (glyc) objectives. Images were then processed using ImageJ (Schneider et al., 2012).

STED imaging was performed with a TCS SP8 STED 3X FALCON confocal head (Leica Microsystems, Germany) mounted on an inverted microscope (DMi8; Leica Microsystems, Germany). For imaging, a 405 nm diode and a white light laser were used as excitation sources for DAPI, ExoGlow-RNA EV (SBI, USA), Alexa Flour 594 (ThermoFisher, USA), and ATTO 647N (ATTO-TEC GmbH, Germany) (405 nm, 488 nm, 575 nm, 644 nm lasers lines respectively). Single photons were collected through a 100×/1.4 NA oil-immersion objective and detected on Hybrid Detectors (HyD) (Leica Microsystems) with a 420 – 500 nm, 500 – 560 nm, 590 – 670 nm, 660 – 720 nm spectral detection window for DAPI, ExoGlow-RNA EV, Alexa Flour 594, and ATTO 647N detection, respectively. For depletion, a 775 nm pulsed laser was used for Alexa Fluor 594 and ATTO 647N, whereas a 660 continuous wave laser was used for depletion of ExoGlow-RNA EV. DAPI was not depleted and only imaged with confocal resolution. The image size was set to 1024 × 1024 pixels and a 5-fold zoom factor was applied, giving a pixel size of 0.023 μm and an image size of 23,25 × 23,25 μm. For FLIM, the white light laser delivered 80 MHz repetition rate. Arrival time of single photons was measured with the included FALCON module and 8 frames were acquired at a scanning speed of 200 Hz. Recordings were done sequentially for each dye to avoid cross-talk. Raw STED images were further processed with the τ-STED module of LAS X software (Leica Microsystems, Germany) increasing further the resolution thanks to the lifetime information recorded.

For the live imaging of neuron exocytosis, young neurons (1-2 weeks) from two control NPC lines and two patient NPC lines were transfected with the plasmid pCMV-Sport6-CD63-pHluorin (A gift from DM Pegtel (Addgene plasmid # 130901 ; http://n2t.net/addgene:130901 ; RRID:Addgene_130901) using the Lipofectamine™ 3000 Transfection Reagent (ThermoFisher Scientific, USA) as instructed in the protocol. 48h following the transfection, cells were imaged using a Leica TIRF system and a 100x/1.47 NA objective as previously described (Verweij et al., 2018). The videos obtained were analyzed using the AMvBE (Analyzer of Multivesicular Body Exocytosis) macro previously developed (Verweij et al., 2018) for ImageJ (Schneider et al., 2012).

### Proteomic analysis

#### -Sample preparation for mass spectrometry

Purified EVs, collected from the conditioned media of 20-30 COs in culture, and IVs, isolated from a pool of 5-7 COs, were lysed in RIPA buffer (150mM NaCl, 50mM Tris pH8, 0.1% DOC, 0.1% SDS, 0.1% NP40). 10 ug of protein for each sample was subjected to the modified FASP protocol (Wiśniewski et al., 2009). Briefly, the protein extract was loaded onto the centrifugal filter CO10 kDa (Merck Millipore, Darmstadt, Germany), and detergent were removed by washing five times with 8M Urea (Merck, Darmstadt, Germany) 50mM Tris (Sigma-Aldrich, USA) buffer. Proteins were reduced by adding 5mM dithiothreitol (DTT) (Bio-Rad, Canada) at 37degrees C for 1 hour in the dark. To remove the excess of DTT, the protein sample was washed three times with 8M Urea, 50mM Tris. Subsequently protein thiol groups were blocked with 10mM iodoacetamide (Sigma-Aldrich, USA) at RT for 45 min. Before proceeding with the enzymatic digestion, urea was removed by washing the protein suspension three times with 50mM ammonium bicarbonate (Sigma-Aldrich, Spain). Proteins were digested first by Lys-C (Promega, USA) at RT for 2 hours, then by trypsin (Premium Grade, MS Approved, SERVA, Heidelberg, Germany) at RT, overnight, both enzymes were added at an enzyme-protein ratio of 1:50 (w/w). Peptides were recovered by centrifugation followed by two additional washes with 50mM ammonium bicarbonate and 0.5M NaCl (Sigma-Aldrich, Swisserland). The two filtrates were combined, the recovered peptides were lyophilized under vacuum. Dried tryptic peptides were desalted using C18-tips (Thermo Scientific, Pierce, USA), following the manufacture instructions. Briefly, the peptides dissolved in 0.1%(v/v) formic acid (Thermo scientific, USA) were loaded onto the C18-tip and washed 10 times with 0.1 % (v/v) formic acid, subsequently the peptides were eluted by 95% (v/v) acetonitrile (Merck, Darmstadt, Germany), 0.1% (v/v) formic acid. The desalted peptides were lyophilized under vacuum. The purified peptides were reconstituted in 0.1% (v/v) formic acid for LC-MS/MS analysis.

#### -MS data acquisition

Desalted peptides were loaded onto a 25 cm, 75 µm ID C18 column with integrated nanospray emitter (Odyssey/Aurora, ionopticks, Melbourne) via the autosampler of the Thermo Easy-nLC 1000 (Thermo Fisher Scientific) at 60 °C. Eluting peptides were directly sprayed onto the timsTOF Pro (Bruker Daltonics). Peptides were loaded in buffer A (0.1% (v/v) formic acid) at 400 nl/min and percentage of buffer B (80% acetonitril, 0.1% formic acid) was ramped from 5% to 25% over 90 minutes followed by a ramp to 35% over 30 minutes then 58% over the next 5 minutes, 95% over the next 5 minutes and maintained at 95% for another 5 minutes. Data acquisition on the timsTOF Pro was performed using timsControl. The mass spectrometer was operated in data-dependent PASEF mode with one survey TIMS-MS and ten PASEF MS/MS scans per acquisition cycle. Analysis was performed in a mass scan range from 100-1700 m/z and an ion mobility range from 1/K0 = 0.85 Vs cm-2 to 1.30 Vs cm-2 using equal ion accumulation and ramp time in the dual TIMS analyzer of 100 ms each at a spectra rate of 9.43 Hz. Suitable precursor ions for MS/MS analysis were isolated in a window of 2 Th for m/z < 700 and 3 Th for m/z > 700 by rapidly switching the quadrupole position in sync with the elution of precursors from the TIMS device. The collision energy was lowered as a function of ion mobility, starting from 45 eV for 1/K0 = 1.3 Vs cm-2 to 27eV for 0.85 Vs cm-2. Collision energies were interpolated linear between these two 1/K0 values and kept constant above or below these base points. Singly charged precursor ions were excluded with a polygon filter mask and further m/z and ion mobility information was used for ‘dynamic exclusion’ to avoid re-sequencing of precursors that reached a ‘target value’ of 14500 a.u. The ion mobility dimension was calibrated linearly using three ions from the Agilent ESI LC/MS tuning mix (m/z, 1/K0: 622.0289, 0.9848 Vs cm-2; 922.0097 Vs cm-2, 1.1895 Vs cm-2; 1221.9906 Vs cm-2, 1.3820 Vs cm-2).

#### -Raw data analysis of MS measurements

Raw data were processed using the MaxQuant computational platform (version 1.6.17.0) (Tyanova et al., 2016) with standard settings applied for ion mobility data (Prianichnikov et al., 2020). Shortly, the peak list was searched against the Uniprot database of Human database (75069 entries, downloaded in July 2020) with an allowed precursor mass deviation of 10 ppm and an allowed fragment mass deviation of 20 ppm. MaxQuant by default enables individual peptide mass tolerances, which was used in the search. Cysteine carbamidomethylation was set as static modification, and methionine oxidation, deamidation and N-terminal acetylation as variable modifications. The match-between-run option was enabled, and proteins were quantified across samples using the label-free quantification algorithm in MaxQuant generating label-free quantification (LFQ) intensities.

#### -Bioinformatic analysis

For the proteomic characterization in IVs and EVs, 7486 proteins were quantified. Proteins that were consistently detected in 2 of the 3 technical replicates per each condition were retained. Downstream analysis was performed using R. The LFQs values were log2-transformed. Missing values were imputed using the R package DEP (version 1.15.0) and replaced by random values of a left-shifted Gaussian distribution (shift of 1.8 units of the standard deviation and a width of 0.3). Differentially expression (DE) analysis was performed on the imputed data using Student’s t-Test. Proteins with log2 fold change values (log2FC) ≥ 1 and ≤ -1 and with an FDR-corrected q-value < 0.05 were considered as differentially expressed. The gene-disease associations analysis was performed using DisGeNET (https://www.disgenet.org/search) (Piñero et al., 2017).

### Validation of proteomic results with automated Western blot

EVs samples were collected from independent new cultures of COs and cells. 1 μg of proteins were loaded on automated western blot system (Proteinsimple WES, https://www.proteinsimple.com) with 12-230 kDa (SM-W004) or 66-440 kDa (SM-W006) Jess/Wess Separation Modules according to the molecular weight of the analyzed protein. All the antibodies were diluted 1:50 (Table 2). Fig. S1F, S2J and S3A show the protein simple profiles and relative quantifications performed using ImageJ Software (https://imagej.nih.gov/).

**Table 2.**
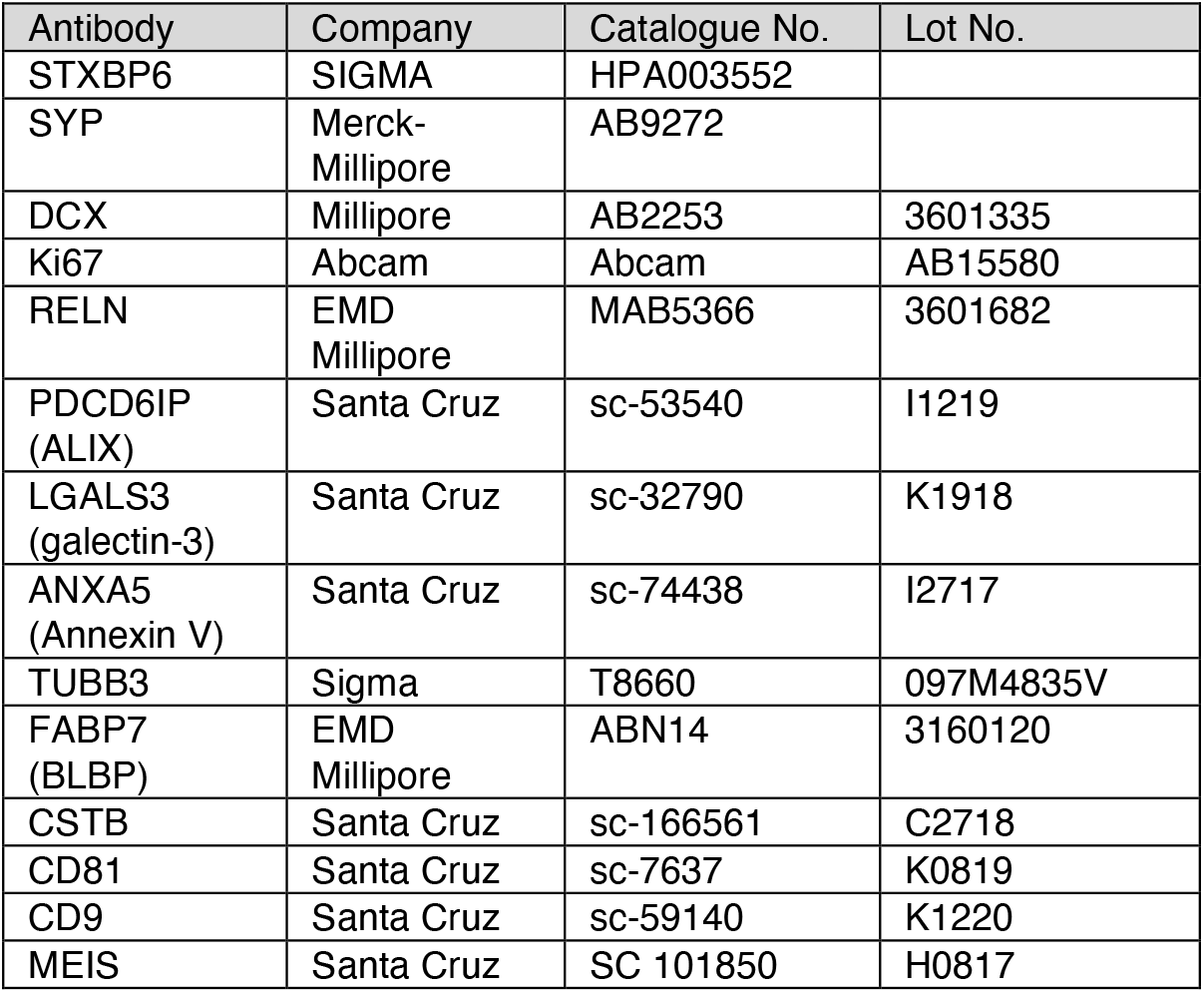
Western blot validation antibodies

### Single-cell RNA-sequencing

Single-cell dissociation was performed on five 60 days old-patterned spheroids randomly selected for each pattern condition. Single cells were dissociated using StemPro Accutase Cell Dissociation Reagent (Life Technologies), filtered through 30 uM and 20 uM filters (Miltenyi Biotec) and cleaned of debris using a Percoll (Sigma, P1644) gradient. Single cells were resuspended in ice-cold Phosphate-Buffered Saline (PBS) supplemented with 0.04% Bovine Serum Albumin at a concentration of 1000 cells per ul. Single cells were loaded onto a Chromium Next GEM Single Cell 3ʹ chip (Chromium Next GEM Chip G Single Cell Kit, 16 rxns 10XGenomics PN-1000127) with the Chromium Next GEM Single Cell 3ʹ GEM, Library & Gel Bead Kit v3.1 (Chromium Next GEM Single Cell 3ʹ GEM, Library & Gel Bead Kit v3.1, 4 rxns 10xGenomics PN-1000128) and cDNA libraries were generated with the Single Index Kit T Set A, 96 rxns (10xGenomics PN-1000213) according to the manufacturer’s instructions. Libraries were sequenced using Illumina NovaSeq6000 in 28/8/91bp mode (SP flowcell), quality control and UMI counting were performed by the Max-Planck für molekulare Genetik (Germany). Downstream analysis was performed using the R package Seurat (version 3.2). Cells with more than 2,500 or less than 200 detected genes or with mitochondrial content higher that 10% were excluded as well as genes that were not expressed in at least three cells. Normalization of gene expression was done using a global-scaling normalization method (“LogNormalize”, scale.factor = 10000) and the 2000 most variable genes were selected (selection method, “vst”) and scaled (mean = 0 and variance = 1 for each gene) before principal component analysis. The “FindNeighbors” and “FindClusters” functions were used for clustering with resolution of 0.5 and UMAP for visualization. Clusters were grouped based of the expression of known marker genes and differentially expressed gene identified with the “FindAllMarkers” function.

### Bulk-RNA-sequencing

RNA-seq was performed on 10ng of total RNA collected from 3 independent wells of NPCs from a 24well plate. NPCs were not treated with EVS or treated for 12h with EVs collected by ultracentrifugation from 25 ml of conditioned medium collected from 28 to 37 days in culture COs (control ventral, EPM1 ventral, control dorsal and EPM1 dorsal COs). NPCs were lysed in 1ml Trizol(Qiagen)/well and RNA was isolated employing RNA Clean & Concentrator kit (Zymo Research) including digestion of remaining genomic DNA according to producer’s guidelines. RNA was further processed according to (Cernilogar et al., 2019). Briefly, cDNA synthesis was performed with SMART-Seq v4 Ultra Low Input RNA Kit (Clontech cat. 634888) according to the manufacturer’s instruction. cDNA was fragmented to an average size of 200–500 bp in a Covaris S220 device (5 min; 4°C; PP 175; DF 10; CB 200). Fragmented cDNA was used as input for library preparation with MicroPlex Library Preparation Kit v2 (Diagenode, cat. C05010012) and processed according to the manufacturer’s instruction. Libraries were quality controlled by Qubit and Agilent DNA Bioanalyzer analysis. Deep sequencing was performed on a HiSeq 1500 system according to the standard Illumina protocol for 50 bp paired-end reads with v3 sequencing reagents.

### RNAseq analysis

Paired end reads were aligned to the human genome version GRCh38 using STAR v2.6.1d (Dobin et al., 2013) with default options “--runThreadN 32 --quantMode TranscriptomeSAM GeneCounts --outSAMtype BAM SortedByCoordinate”. Reads-per-gene counts were imported in R v4.1.0. Bioconductor package DESeq2 v1.32.0 (Love et al., 2014) was used for differential expression analysis. Only genes with read counts>1 were considered. Significantly changed genes were determined through pairwise comparisons using the DESeq2 results function (log2 fold change threshold=1, adjusted p-value <0.05). Heatmaps with differentially expressed genes were plotted with pheatmap v1.0.12 and RColorBrewer v1.1-2 using rlog-normalized expression values.

### Enrichment analysis

GO term analysis of differentially expressed proteins was tested using the FUMA algorithm (Watanabe et al., 2017) by inserting the DE protein lists into the GENE2FUNC software (FDR<0.05) (https://fuma.ctglab.nl/) or with STRING (https://string-db.org).

## Acknowledgments

we thank E.Frenna, I.Miinalainen, C.Kyrousi, V.Pravata, M.Ködel, A.Krontira, P.Lopez, C.Cruceanu, G.Berto, C.Turk, S.Zünd, A.C.Ayo-Martin, S.Dietzel and P.Gressens for technical help and critical discussion.

## Author contributions

Conceptualization: SC, RDG. Methodology: RDG, FP, AF, FDM, GM, PK, SM, LC, CG, MWP, MGP, ZB. Investigation: SC, RDG, FP, AF, FDM, NB, FC. Visualization: SC, FP, AF, SM, BP, MGP. Funding acquisition: SC. Supervision: SC, RDG, GM, DJ. Writing – original draft: SC; review & editing: SC, RDG, FP, AF.

---

**Supplementary Fig. S1.**
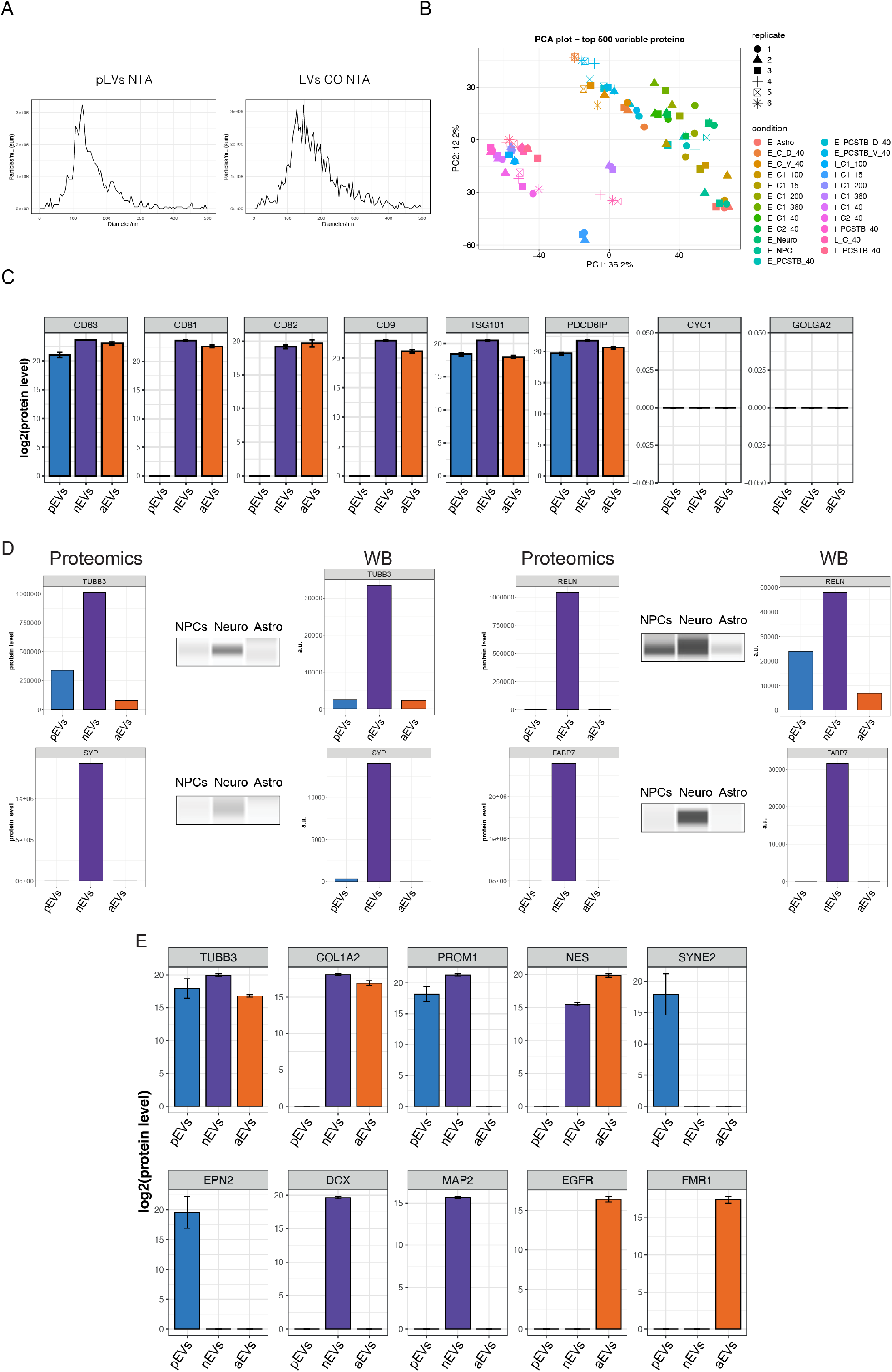
(A) NTA plot of EVs isolated by differential ultracentrifugation from NPCs (pEVs) and COs (EVs CO). (B) Principal component Analysis (PCA) plot of protein samples from different CO and cell type vesicles, based on LFQ intensity of quantified proteins. All the replicates are represented. (C) Bar plots showing positive and negative EV marker expression in pEVs, nEVs, and aEVs. Data are represented as mean ± SD of 3 technical replicates. EVs were collected from the conditioned media of 15cm petri dishes containing 20-30 COs. (D) Validation of proteomics (left) by Western blot analysis (middle) and relative quantifications (arbitrary units, a.u.) (right) on protein extracts from EVs from different cell types. EVs were collected from the conditioned media of 3-5 different wells of cells in culture. (E) Bar plots showing cell-type specific markers in pEVs, nEVs, and aEVs. Data are represented as mean ± SD of 3 technical replicates. EVs were collected from the conditioned media of 3-5 different wells of cells in culture.

**Supplementary Fig. S2.**
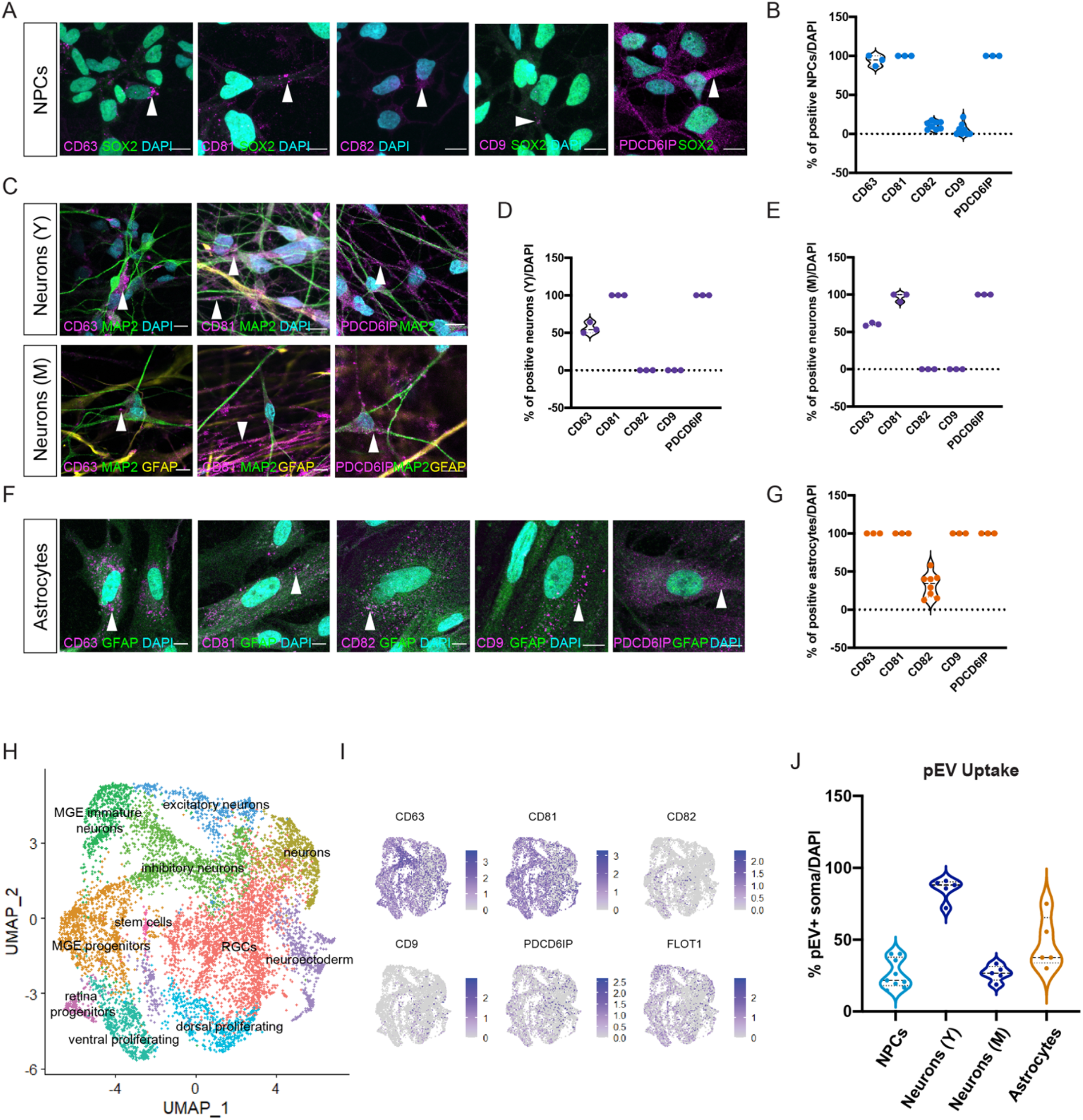
(A) CD63, CD81, CD82, CD9, PDCD6IP immunostaining in NPCs (SOX2+, green) (arrow heads). DAPI, cyan. Scale bar: 10 μm. (B) Quantification of CD63+, CD81+, CD82+, CD9+ and PDCD6IP+ NPCs. Violin plots show median and interquartile range. Every dot refers to a field of view, n= 3-8 per condition. (C) CD63, CD81, PDCD6IP immunostaining in young neurons (Y) (MAP2+, green) (top) and mature neurons(M) (MAP2+, green and GFAP+, yellow) (bottom) (arrow heads). DAPI, cyan. Scale bar: 10 μm. (D) Quantification of CD63+, CD81+ and PDCD6IP+ neurons (Y). Violin plots show median and interquartile range. Every dot refers to a field of view, n= 3 per condition. (E) Quantification of CD63+, CD81+ and PDCD6IP+ neurons (M). Violin plots show median and interquartile range. Every dot refers to a field of view, n= 3 per condition. (F) CD63, CD81, CD82, CD9, PDCD6IP immunostaining in astrocytes (GFAP+, green) (arrow heads). DAPI, cyan. Scale bar: 10 μm. (G) Quantification of CD63+, CD81+, CD82+, CD9+ and PDCD6IP+ astrocytes. Violin plots show median and interquartile range. Every dot refers to a field of view, n=3-8 per condition. (H) UMAP plot of scRNA-seq clusters of vCOs and dCOs. n= 3 per condition. (I) Feature plots showing the expression of vCO and dCO EV markers (as shown in the panel). n= 3 per condition. (J) Quantification of pEV uptake by NPCs, neurons (Y), meurons (M), and astrocytes. Violin plots show median and interquartile range. Every dot refers to a field of view, n= 5-9 per condition.

**Supplementary Fig. S3.**
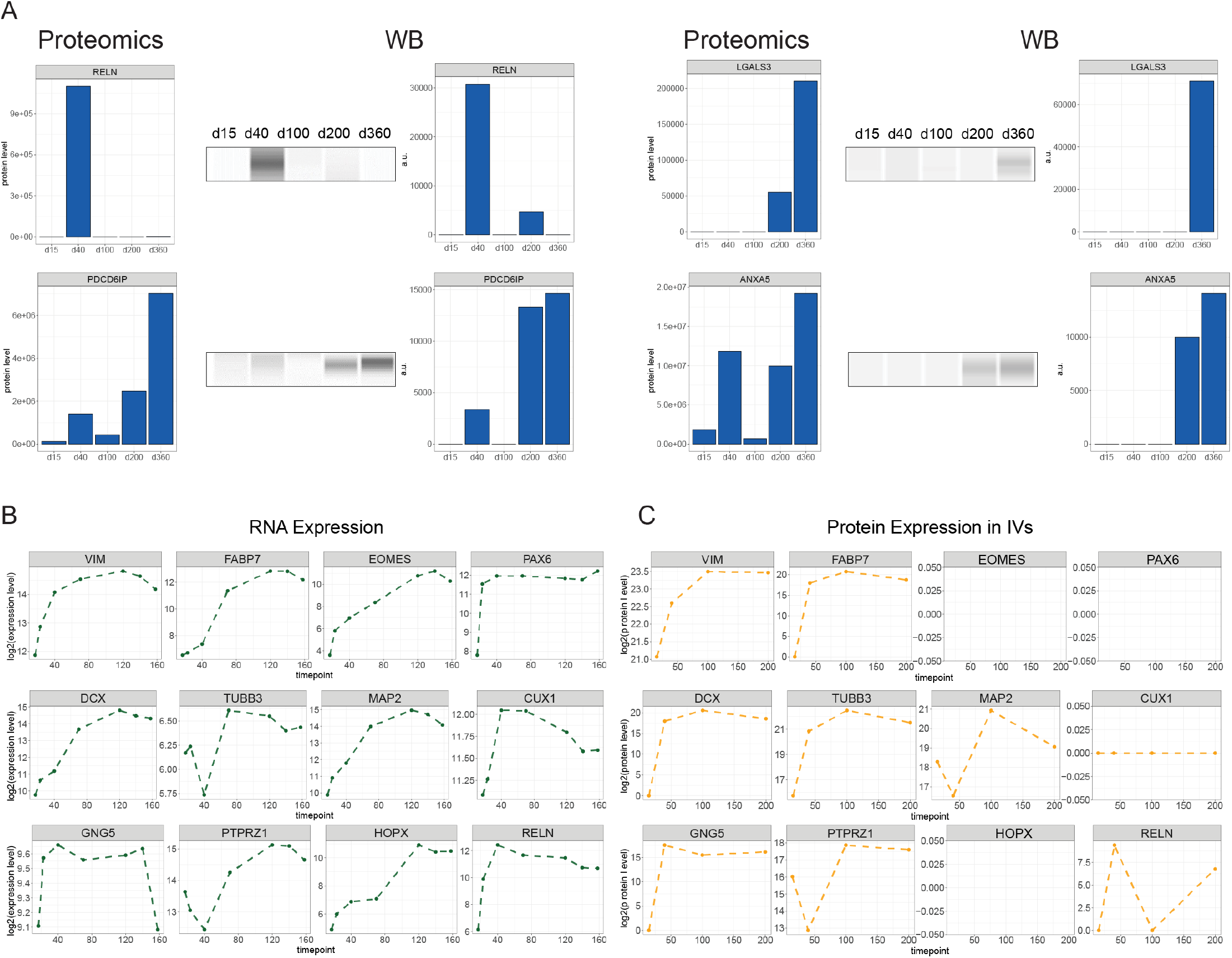
(A) Validation of proteomics (left) by Western blot analysis (middle) and relative quantifications (arbitrary units, a.u.) (right) on protein extracts from EVs from COs at different developmental stages (d0-d360). EVs were collected from the conditioned media of 15cm petri dishes containing 20-30 COs. (B) Temporal trajectories of RNA expression of developmental markers in COs (bulk RNA-seq (Cruceanu et al., 2022)). (C) Temporal trajectories of protein levels of developmental markers in IVs from COs. n= 3 per condition.

**Supplementary Fig. S4.**
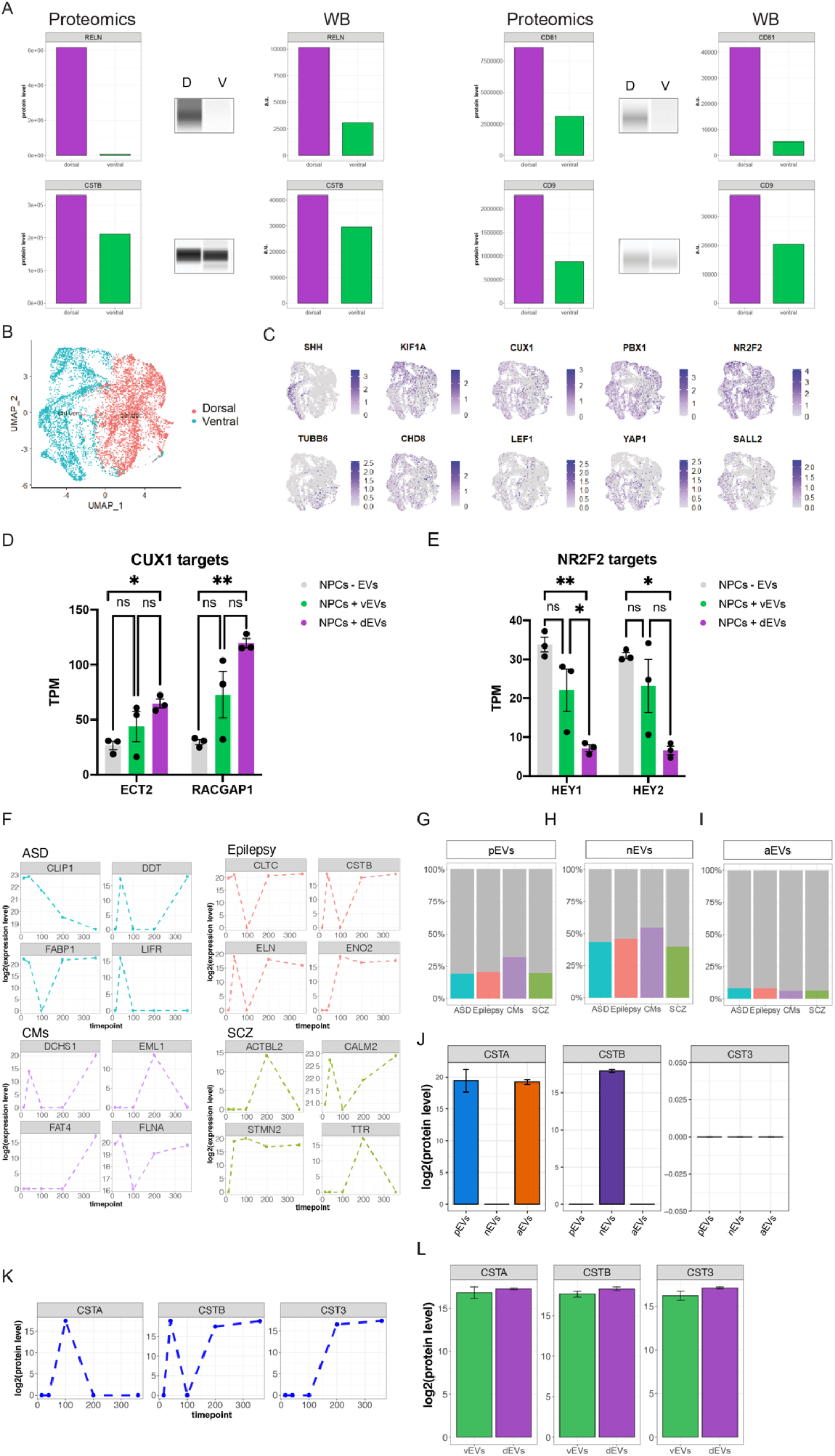
(A)Validation of proteomics (left) by Western blot analysis (middle) and relative quantifications (arbitrary units, a.u.) (right) on protein extracts from EVs in brain-region-specific CO EVs. EVs were collected from the conditioned media of 15cm petri dishes containing 20-30 COs. (B) UMAP plot of scRNA-seq clusters of vCOs and dCOs. N= 3 per condition. (C) Feature plots showing the expression of vCO and dCO EV markers (as shown in the panel). n= 3 per condition. (D) Bar plots showing the expression (TPM, Transcripts Per Million) of CUX1 targets (ECT2 and RACGAP1) in NPCs after treatment with no EVs, vEVs, and dEVs. Data are represented as mean and ± SEm. n= 3 per condition. Statistical significance was based on a one-way analysis of variance (ANOVA), *p<0.05, **p<0.01. (E) Bar plots showing the expression (TPM, Transcripts Per Million) of NR2F2 targets (HEY1 and HEY2) in NPCs after treatment with no EVs, vEVs, and dEVs. Data are represented as mean and ± SEm. n= 3 per condition. Statistical significance was based on a one-way analysis of variance (ANOVA), *p<0.05, **p<0.01. (F) Temporal trajectories of proteins associated to ASD (Autism Spectrum Disorder, cyan), Epilepsy (red), CMs (Cortical Malformations, purple) and SCZ (Schizophrenia, green) in EVs from COs. (G) Bar plots indicating the proportion of proteins associated with neurodevelopmental disorders found in CO EVs with proteins secreted in EVs from NPCs, (H) neurons and (I) astrocytes (https://www.disgenet.org/). Data are represented as mean ± SD of technical replicates. n= 3 per condition. (J) Bar plots of the protein levels of the cystatin family (CSTA, CSTB, CST3) in pEVs, nEVs, and aEVs. Data are represented as mean ± SD of 3 technical replicates. EVs were collected from the conditioned media of 3-5 different wells of cells in culture. (K) Developmental trajectory of protein levels of the cystatin family (CSTA, CSTB, CST3) in EVs. (L) Bar plots of the protein levels of the cystatin family (CSTA, CSTB, CST3) in vEVs and dEVs. Data are represented as mean ± SD of 3 technical replicates. EVs were collected from the conditioned media of 3-5 different wells of cells in culture.

**Supplementary Fig. S5.**
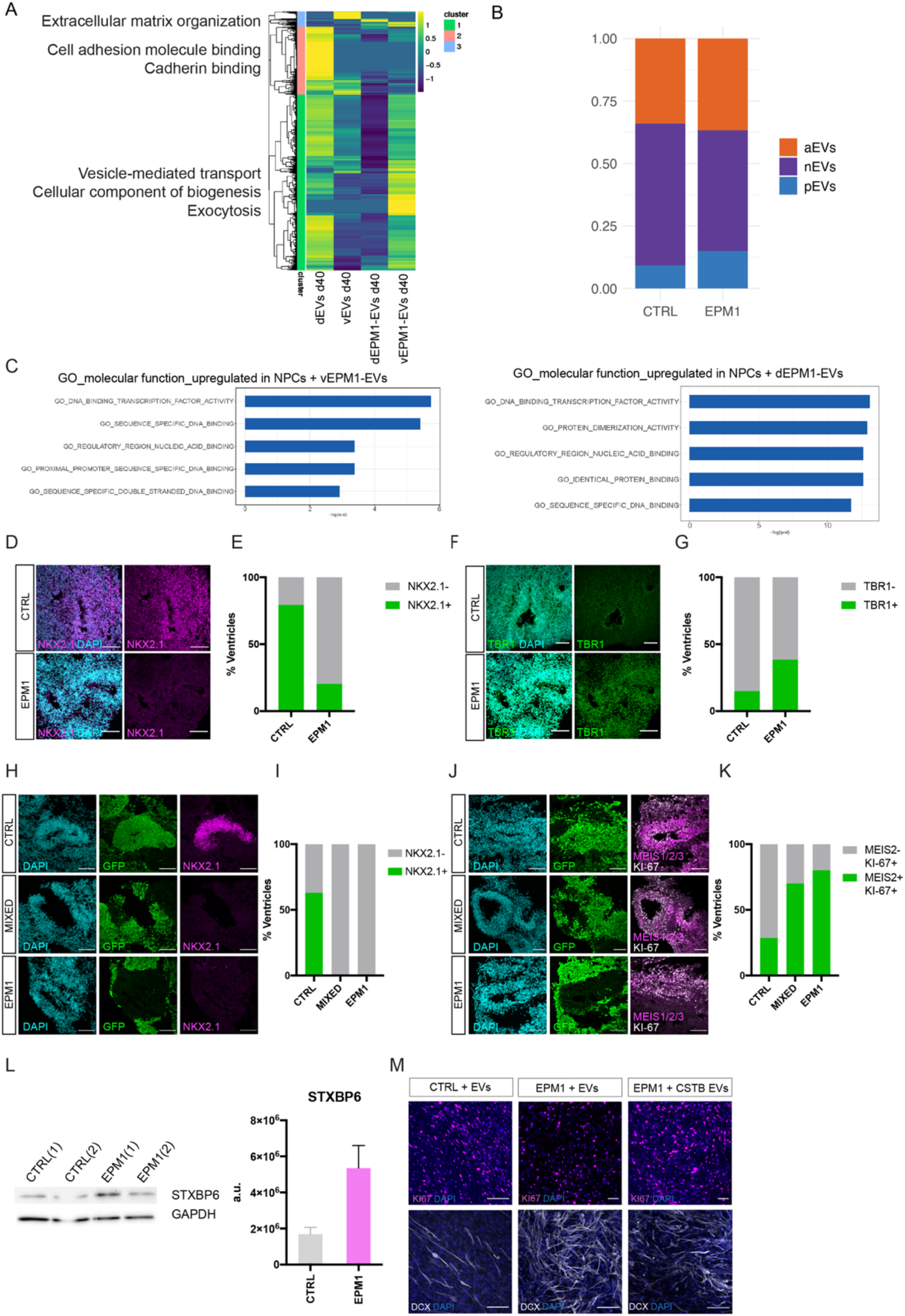
(A) Heatmap showing hierarchical clusters of vEV, dEV, vEPM1-EV and dEPM1-EV proteins. EVs were collected from the conditioned media of 15cm petri dishes containing 20-30 COs. EV proteomic analysis was performed on EPM1 patient-derived COs from two different patients (pooled together for this analysis). (B) Bar plot comparing EPM1-EV DE proteins with the protein content of pEVs, nEVs and aEVs. (C) GO enrichments for up-regulated genes in NPCs after acute treatment with vEVs and vEPM1EVs (left), and with dEVs and dEPM1-EVs (right). GOs for molecular function are reported. (D) Immunostaining of vCOs and vEPM1-COs at 30d for NKX2.1+ ventral progenitors (magenta) and DAPI (light blue). Scale bar: 100 µm. (E) Quantification of NKX2.1+ventricles in vCOs and EPM1-vCOs. 63 ventricles and 69 ventricles were counted in vCOs and EPM1-vCOs respectively. Statistical significance was based on the binomial test *p<0.05, **p<0.01, ****p<0.0001. (F) Immunostaining of vCOs and EPM1-vCOs at 30d for TBR1 dorsal neurons (green) and DAPI (light blue). Scale bar: 100 µm. (G) Quantification of TBR1+ventricles in vCOs and EPM1-vCOs. 67 ventricles and 52 ventricles were counted in vCOs and EPM1-vCOs respectively. Statistical significance was based on the binomial test *p<0.05, **p<0.01, ****p<0.0001. (H) Immunostaining of GFP+vCOs, Hybrid-vCOs and EPM1-vCOs at 70d for NKX2.1 ventral progenitors (magenta), GFP control cells (green) and DAPI (light blue) Scale bar: 100 µm. (I) Quantification of NKX2.1+ventricles in GFP+vCOs, Hybrid-vCOs and EPM1-vCOs. 19 GFP+ ventricles, 10 GFP-ventricles and 14 hybrid ventricles were counted. Statistical significance was based on the binomial test *p<0.05, **p<0.01, ****p<0.0001. (J) Immunostaining of GFP+vCOs, Hybrid-vCOs and EPM1-vCOs at 70d for proliferating MEIS2-KI67 ventral progenitors (magenta), GFP control cells (green) and DAPI (light blue) Scale bar: 100 µm. (K) Quantification of proliferating MEIS2-KI67+ventricles in GFP+vCOs, Hybrid-vCOs and EPM1-vCOs. 9 GFP+ ventricles, 10 GFP-ventricles and 10 hybrid ventricles were counted. Statistical significance was based on the binomial test *p<0.05, **p<0.01, ****p<0.0001. (L) Western Blot analysis of STXBP6 and GAPDH on protein extracts from CTRL and EPM1 CO (right), and relative quantifications (arbitrary units, a.u.) (left). Total cell lysates were obtained from a pool of 5-7 COs. (M) Representative immunostaining showing KI67+ (magenta) and DCX+ (white)NPCs derived from control and EPM1 COs after EV treatment. DAPI, cyan. Scale bar: 50 μm.

